# DEXOM: Diversity-based enumeration of optimal context-specific metabolic networks

**DOI:** 10.1101/2020.07.17.208918

**Authors:** Pablo Rodríguez-Mier, Nathalie Poupin, Carlo de Blasio, Laurent Le Cam, Fabien Jourdan

## Abstract

The correct identification of metabolic activity in tissues or cells under different environmental or genetic conditions can be extremely elusive due to mechanisms such as post-transcriptional modification of enzymes or different rates in protein degradation, making difficult to perform predictions on the basis of gene expression alone. Context-specific metabolic network reconstruction can overcome these limitations by leveraging the integration of multi-omics data into genome-scale metabolic networks (GSMN). Using the experimental information, context-specific models are reconstructed by extracting from the GSMN the sub-network most consistent with the data, subject to biochemical constraints. One advantage is that these context-specific models have more predictive power since they are tailored to the specific organism and condition, containing only the reactions predicted to be active in such context. A major limitation of this approach is that the available information does not generally allow for an unambiguous characterization of the corresponding optimal metabolic sub-network, i.e., there are usually many different sub-network that optimally fit the experimental data. This set of optimal networks represent alternative explanations of the possible metabolic state. Ignoring the set of possible solutions reduces the ability to obtain relevant information about the metabolism and may bias the interpretation of the true metabolic state. In this work, we formalize the problem of enumeration of optimal metabolic networks, we implement a set of techniques that can be used to enumerate optimal networks, and we introduce `DEXOM`, a novel strategy for diversity-based extraction of optimal metabolic networks. Instead of enumerating the whole space of optimal metabolic networks, which can be computationally intractable, `DEXOM` samples solutions from the set of optimal metabolic sub-networks maximizing diversity in order to obtain a good representation of the possible metabolic state. We evaluate the solution diversity of the different techniques using simulated and real datasets, and we show how this method can be used to improve in-silico gene essentiality predictions in *Saccharomyces Cerevisiae* using diversity-based metabolic network ensembles. Both the code and the data used for this research are publicly available on GitHub^1^.

## Introduction

Metabolism and its regulation is an ensemble of intrincated and tightly coordinated processes involving hundreds to thousands of enzymes, reactions, metabolites and genes, whose interactions define complex networks that are unique for each species. This complexity grants organisms the flexibility to adapt their energetic functions and growth requirements to a wide variety of conditions. Changes in nutrient availability, conditions of cellular stress, or any other change in the environment can induce a rapid metabolic reprogramming of cells, rewiring their metabolism to adjust to the requirements of the new situation. Dysfunction of these mechanisms play a central role in the development of many diseases, but most notably in cancer, where cancer cells exploit metabolic reprogramming on their own benefit [1] to sustain a rapid proliferation rate and survive in conditions of hypoxia, nutrient depletion, or even develop therapy resistance [2]. Being able to accurately detect these changes or deregulations in metabolism would be beneficial not only for a better understanding of biological systems but to develop more targeted therapies and treatments for many diseases [3–5].

One of the reasons why this task remains elusive is the complexity of the multiple processes that participate in the regulation of the metabolism [6]. More specifically, post-transcriptional control of mRNA, post-translational modifications of enzymes, as well as biochemical constraints —including for example the laws for mass and charge conservation, cell growth requirements, biomass composition and nutrient availability— make the identification of which pathways are altered between conditions very complicated by the mere observation of changes in gene expression or changes in metabolite concentrations. Instead, integrating and analyzing together all those different levels of information is key to improve the predictive models and to provide a more accurate mechanistic view of the system under study.

Genome-scale metabolic networks (GSMN) are suitable computational models for the integration of these multiple levels of knowledge. These models are automatically built and manually curated networks that encode all reactions with their stoichiometric coefficients, metabolites, enzymes, gene annotations and biochemical constraints that are known for an organism. GSMNs are generic models of an organism, independent of the type of tissue, cell or condition. In order to generate more accurate models for specific tissues or conditions, experimental data such as gene or protein expression can be integrated on top of GSMNs using context-specific network reconstruction methods. Taking into account the different levels of expression of genes between conditions, a sub-network from the GSMN is extracted by finding a steady-state flux most consistent with the experimental data. This process allows the generation of metabolic networks specifically tailored to the condition, to highlight for example differences in metabolism between tissues [7–9] or to discover novel drug targets or essential genes in cancer cells [10–12].

Several methods were proposed in the literature to automatically reconstruct context-specific metabolic networks from gene or protein expression [7–9,13–17]. This process is done by solving an optimization problem to find the sub-network from the GSMN that maximizes the agreement with the experimental data. This agreement is defined in different ways: some methods such as [7, 15] use data to classify reactions into reactions associated to highly expressed enzymes (or core reactions) or reactions associated to lowly expressed enzymes, whereas others [8, 14] assign different scores (weights) to reactions based on data and other experimental evidence. The optimization problem is then defined as that of finding the sub-networks that can carry a steady-state flux through the reactions that maximize the overall score. However, a major limitation that is frequently neglected is that the available information is usually not sufficient to fully and unambiguously characterize the corresponding metabolic sub-network for a given condition. Instead a range of different optimal metabolic sub-networks may exist, offering slightly different hypotheses of the possible metabolic state. Ignoring this variability can not only lead to incorrect or incomplete explanations of the biological experiment, but also causes valuable information to be lost that could be used to improve predictions. Although this limitation is starting to be acknowledged [18], there is still a lack of studies that provide methods to analyze and explore the optimal space of alternative networks.

One of the initial works that exploits the idea of multiple context-specific networks to improve predictions is `EXAMO` [19]. In this work, authors perform an enumeration of optimal metabolic networks using `iMAT` [7]. The enumeration is done using the same strategy proposed in `iMAT` for assigning confidence scores to reactions, followed by a post-processing step using the `MBA` [13] algorithm to generate a single consensus network including the reactions predicted to be active. A similar strategy was applied by Poupin et al. [18], but instead of generating a single consensus network, the whole set of networks derived by forcing fluxes through each reaction in the model is preserved as alternative hypotheses of the metabolic state. This enables a better characterization of the metabolic shifts that occur during hepatic differentiation.

The procedure of generating alternative networks by forcing or blocking flux through each reaction has however some limitations. First, it can generate many duplicated solutions. For example, if there exist only one optimal metabolic network with a linear pathway of 10 reactions, forcing the activation of each reaction in the linear pathway will generate always the same optimal solution, wasting computational resources. Second, it cannot recover the whole set of possible optimal metabolic networks, as not all possible combinations of reactions are tested. Third, there is no guarantee that the solution set is representative and diverse of the full space of possible networks. A simple brute force algorithm that could be used to prevent this would be to test every possible combination between variable reactions. However, this approach does not scale as the number of problems to solve grows exponentially with the number of variable reactions. As an alternative to this approach, authors in [20] present a strategy to generate alternative metabolic networks. Of particular interest is their `CorEx` algorithm, which in a similar fashion as `Fastcore` method [15], calculates the smallest flux-consistent

sub-network that preserve the reactions in the core set, but solving the problem exactly instead of the LP-based fast approximations used in `Fastcore`. `CorEx` also incorporates a mechanism to enumerate optimal networks by maximizing the dissimilarity with the previously found solution, a process that can be repeated iteratively to discover new optimal networks. However, without a mechanism that prevents the generation of duplicated solutions, the enumeration process can get stuck in a small region in the space of optimal solutions. Some issues still remain with this enumeration strategy, mainly regarding its effectiveness to get a representative set of the possible metabolic networks and also how to take advantage of the set of networks to improve predictions more than just only observing the variability in terms of reactions that can appear or not in the different optimal sub-newtorks.

Regarding this last question, it was shown that the use of ensembles of draft networks reconstructed using Gap Filling methods with multiple media conditions and random perturbations can improve flux-based predictions [21]. Although the application is different, predictions using context-specific network reconstruction methods could be also improved using ensembles of optimal metabolic networks, and diversity can play an important role in the quality of the ensemble models.

As a response to the current limitations, in this work we develop and analyze different techniques for enumeration of optimal metabolic networks, and we evaluate how well they perform in terms of diversity of the solutions and predictive capabilities (Fig. 1). Based on this analysis, we introduce `DEXOM`, a novel method for diversity-based exploration of context-specific metabolic networks. Using experimental data for a particular condition and organism, we first construct an initial set of sub-networks by testing single variations of reactions that may or may not be present in the networks without affecting the optimality. This set is then expanded to explore combinations of reactions by sampling new sub-networks from the unknown space of possible optimal sub-networks, maximizing diversity in order to detect as much differences as possible. The sampling process is required to obtain a representative set of solutions that cannot be retrieved by exhaustive search due to a combinatorial explosion problem. We evaluate and compare all the methods in terms of 1) diversity of the solution set, and 2) predictive capabilities of ensembles of optimal networks built with each method for the prediction of essential genes in *Saccharomyces Cerevisiae* under aerobic conditions.

**Fig 1.**
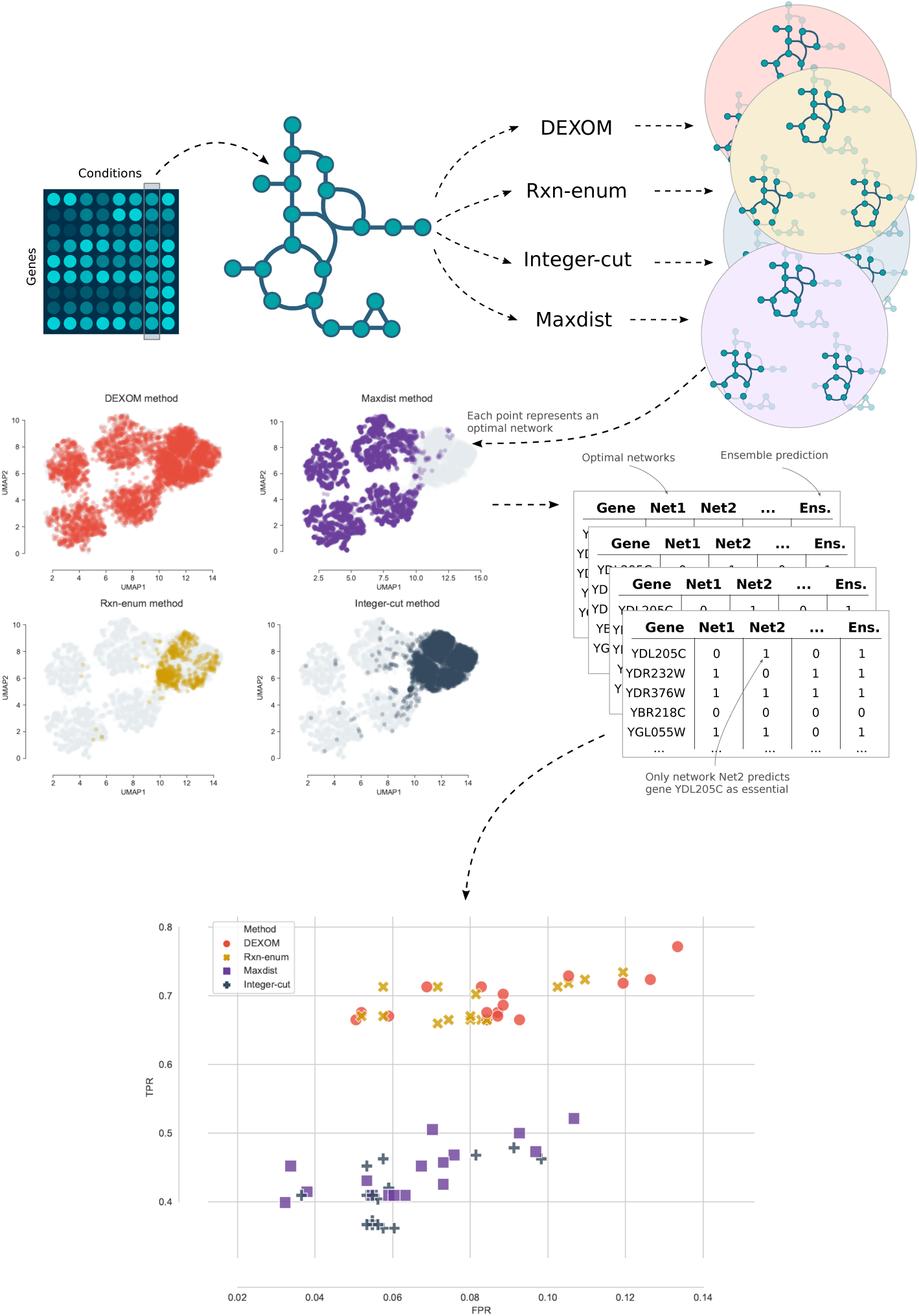
Summary of the approach. The process starts with some experimental data (e.g., gene expression for a given condition and organism) and a GSMN model for the organism. Data is mapped onto the GSMN and the proposed methods are used to generate multiple optimal context-specific metabolic networks. Each set of optimal solutions is compared in terms of diversity, and projected into a 2D embedding to visualize which parts of the space of optimal metabolic networks is explored by each method. The set of optimal networks obtained with each method are used to build ensembles and predict essential genes in yeast.

## Methods

In this section we define the problem of context-specific reconstruction of optimal metabolic networks. We introduce the problem of enumeration of networks, we propose and analyze different strategies for enumeration, and we discuss the advantages and limitations of each approach. Finally we introduce `DEXOM`, our proposed method for context-specific metabolic network reconstruction and enumeration.

### Optimal context-specific metabolic network reconstruction

Here we consider the reconstruction of optimal context-specific metabolic networks as the selection of a subset of reactions from a global genome-scale metabolic network for a particular organism, in a way that maximizes the agreement with experimental data, i.e., reactions in the model with evidence of being active in a given context should be preserved, and reactions with evidence of being inactive should be removed from the model. The selection of this subset of reactions is also subject to flux-based constraints, which constrain the space of possible ways in which those reactions can be selected.

More formally, given:

- *G* = {*R, M, S*}, an initial genome-scale metabolic network *G* for a given model organism, where *R* = {*R*_1_, …, *R*_*n*_} is the set of reactions in the network, *M* = {*M*_1_, …, *M*_*m*_} is the set of metabolites, and *S* is the stoichiometry matrix of size *m* × *n*
- *f* (***x***) : {0, 1}^*n*^ → ℝ, a linear objective function of the form ***c***^***T***^ ***x*** that returns a score for a candidate subset of reactions indexed by a binary vector ***x*** ∈ {0, 1}^*n*^, indicating whether reaction *R*_*i*_ is selected or not, so that the subset of selected reactions from *R* is defined as *R*_*c*_ = {*R*_*i*_ ∈ *R* | *x*_*i*_ = 1, ∀*i* ∈ 1 … *n*}

The goal is to find the binary vector ***x*** (or equivalently the subset *R*_*c*_) such that *f* (***x***) is maximized. Reactions included in the *R*_*c*_ set have to carry a non-zero flux under steady state conditions. This problem can be stated as a Mixed Integer Linear Programming (MILP) problem with the following form:

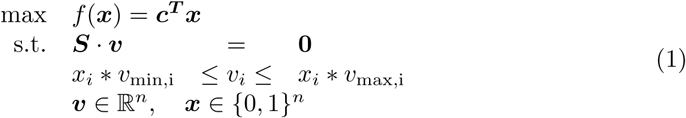

where *x*_*i*_ ∈ {*x*_1_, …, *x*_*n*_} are the binary variables representing if reaction *R*_*i*_ is present or not, *v*_*i*_ ∈ {*v*_1_, …, *v*_*n*_} the variables representing the flux through each reaction *R*_*i*_, and *v*_*min*_ and *v*_*max*_ the lower and upper bounds for the flux through each reaction. Note that what is subject to optimization is the selection of the reactions but not the fluxes. Fluxes are constrained within some bounds ***v***_***min***_ and ***v***_***max***_, and forced to be in steady state (***S*** · ***v*** = 0). Reactions can be included (*x*_*i*_ = 1) only if they can carry some non-zero flux, and reactions not included (*x*_*i*_ = 0) are forced to carry a zero flux. In the following, we shall use this notation to introduce different MILP problems for context-specific reconstruction of metabolic networks.

The objective function *f*(***x***) calculates the agreement between the experimental data and the selected reactions. One common strategy is to divide reactions in two disjoint sets based on experimental evidence, namely reactions associated with highly expressed enzymes (*R*_*H*_ ⊆ *R*) and reactions associated with lowly expressed enzymes (*R*_*L*_ ⊆ *R*), and then defining *f* (***x***) as:

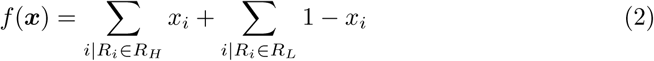

This is the strategy described in `iMAT`, in which the selection of one reaction in *R*_*H*_ or the removal of one reaction in *R*_*L*_ contribute in the same way to the score. Other strategies such as `Fastcore`, enforce the inclusion of all the reactions in *R*_*H*_, and so *f* (***x***) evaluates only the number of selected reactions in *R*_*L*_ to minimize it.

In practice, the binary vector ***x*** is extended to account also for reversible reactions in the *R*_*H*_ set that can be active carrying a negative flux. Also, a tunable parameter *E* corresponding to the minimal amount of flux a reaction has to carry to be considered active is usually included in the optimization problem. In the original `iMAT` formulation, a reaction *R*_*i*_ ∈ *R*_*L*_ which is not selected (which carries no flux) has a value of *x*_*i*_ = 1 representing a match with the experimental data, and so Eq. 2 simplifies to just *f* (***x***) = ∑_*i*_ *x*_*i*_. Full problem specification is described in Eq. 3:

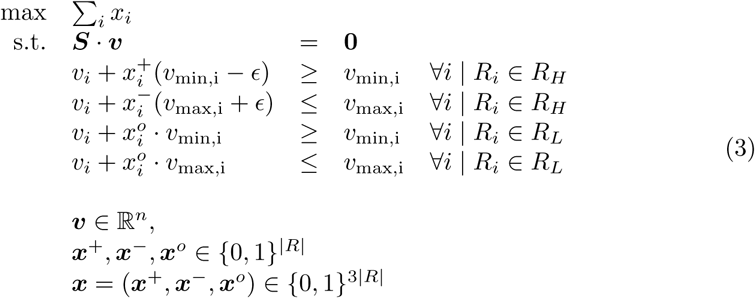

In the following sections, for practical reasons, and without loss of generality, we use the original set of `iMAT` constraints and objective function as the base MILP problem for network enumeration, since: 1) it relies on a MILP formulation, which can be easily adapted to optimally solve different optimization problems and objectives; and 2) the default objective function optimizes a trade-off between the coverage of reactions associated with highly expressed genes and reactions associated with lowly expressed genes, which has been proven in practice a good general strategy that only requires gene expression data. This trade-off introduces flexibility in the optimization process, allowing us to predict that some reactions are not active even though they are associated with highly expressed genes, something important to account for post-transcriptional events.

### The problem of enumerating optimal metabolic networks

The enumeration problem arises naturally in context-specific reconstruction of metabolic networks due to the discrete nature of the selection of reactions.

The enumeration problem can be easily illustrated using the toy example depicted in Fig. 2. The figure shows a toy metabolic network using a Direct Acyclic Graph (DAG) representation with *L* layers of *N* metabolites. Each metabolite *m*_*i,k*_ in layer *L*_*k*_ is connected to each metabolite *m*_*j,k*+1_ in *L*_*k*+1_ by single reactions *R*_*ijk*_ = (*m*_*i,k*_, *m*_*j,k*+1_) with only one substrate and product. The model includes two extra metabolites *m*_*s*_ as a source and *m*_*t*_ as a sink node to centralize the import and export reactions and simplify the model. The number of total metabolites, including *m*_*s*_ and *m*_*t*_ is 2 + *N* · *L*, and the number of total reactions is 2*N* + *N* ^2^ (*L* − 1).

**Fig 2.**
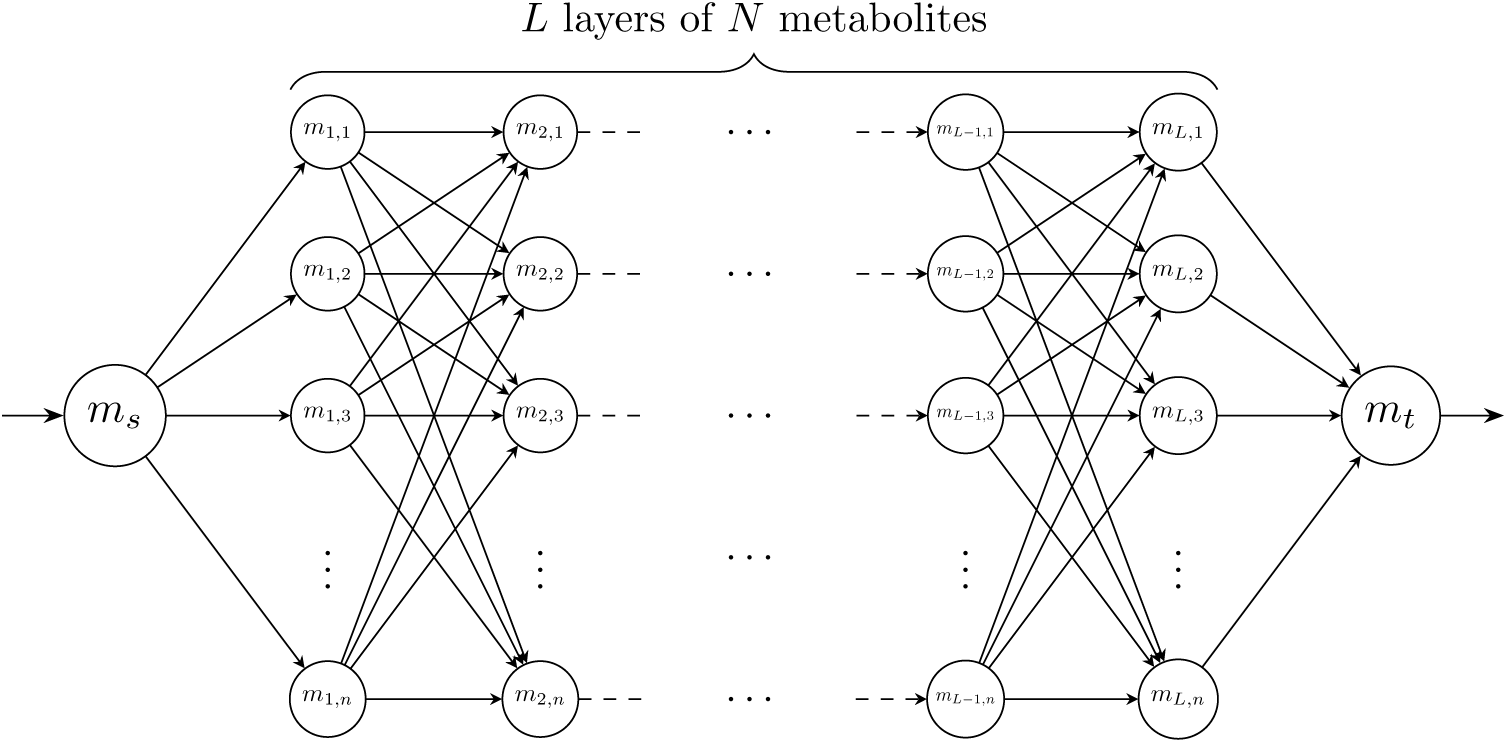
Example of an abstract artificial metabolic network

In this example, we want to extract the context-specific metabolic network, given the following conditions:

- ∑_*i*_|*v*_*i*_| > 0, i.e., there is a non-zero steady state flux from *m*_*s*_ to *m*_*t*_. This is commonly assumed in order to avoid having an empty network.
- *R*_*H*_ = ø, *R*_*L*_ = *R*, i.e., there are no reactions associated to highly expressed enzymes, and all the reactions are associated with lowly expressed enzymes.

It can be seen that a metabolic network with optimal *f* (***x***) in this case is the one that carries flux from *m*_*s*_ to *m*_*t*_ using the minimum number of reactions (since they are all in the *R*_*L*_ set), which corresponds to a shortest path from *m*_*s*_ to *m*_*t*_. Since there are no loops in the network, the shortest length for the path is *L* + 2 (including the path from *m*_*s*_ to *L*_1_ and from *L*_*N*_ to *m*_*t*_). This also implies that there is no single solution, but instead any path from *m*_*s*_ to *m*_*t*_ is an optimal solution, i.e., a context-specific reconstruction network with optimal *f* (***x***) given the previously defined conditions. Since there are *N* different paths to go from any metabolite in layer *L*_*j*_ to any metabolite in layer *L*_*j*+1_, that makes *N*^*L*^ possible optimal networks in this particular example, that is, the number of possible optimal solutions in this example grows exponentially with the number of layers. Note also that since the number of reactions for a fixed number of metabolites grows linearly with the number of layers, the number of possible solutions grows also exponentially with the number of reactions.

This example illustrates that there are instances of the enumeration problem for which the number of optimal solutions grows exponentially with the size of the network. Thus, in general, enumerating the full set of optimal metabolic networks can be impractical, especially considering the size of networks such as Recon 3D [22] with 13,543 reactions, or the recent Human1 network [23] with around 13,000 reactions.

More formally, it can be shown that the enumeration of all optimal metabolic networks is a type of *vertex enumeration problem*. Let *M*_*P*_ be the general MILP problem for context specific reconstruction using a GSMN with *n* reactions and with objective function ***c***^***T***^ ***x*** that we want to maximize, as defined in Eq. 1. Let Ω be the set of all 0/1-vectors representing the feasible solutions for the MILP *M*_*P*_ that satisfy all the constraints defined in Eq. 1. From a geometric point of view, the space of possible networks can be viewed as vertices of the hypercube *C*_*n*_ = {0, 1}^*n*^, and the set of feasible solutions Ω as a subset of vertices of *C*_*n*_, where its convex hull is a 0/1-polytope *P*, that is, *P* = *conv*(Ω). Let *z*^∗^ be the optimal value of *M*_*P*_, i.e., ∀***x*** ∈ Ω, ***c***^***T***^ ***x*** ≤ *z*^∗^. We are interested in the set of all optimal feasible solutions Ω^∗^ := {***x***^**∗**^|***x***^**∗**^ ∈ Ω ∧ ***c***^***T***^***x***^**∗**^ = *z*^∗^}, where *P*^∗^ = *conv*(Ω^∗^) ⊆ *P* is the 0/1-subpolytope of interest in -representation (as the intersection of half spaces defined by all the constraints) from which we want to obtain the 𝒱-representation, that is, the set of vertices as vectors of 0/1 coordinates (the optimal context-specific networks), which is the definition of the *vertex enumeration problem*.

Vertex enumeration [24] is a classical problem in the field of combinatorial optimization for which some specific techniques were proposed [25]. For the special case of 0/1-polytopes [26], some notable approaches are Binary Decision Diagrams [27–29], tree search-based methods [30, 31] and techniques based on branch-and-bound and cutting planes, extensively exploited in commercial solvers such as IBM CPLEX and Gurobi. In fact, as a general enumeration mechanism, these solvers incorporate the concept of a pool of optimal solutions, in which the tree of feasible solutions continues to be explored until a specific number of optimal feasible solutions have been found.

However, as discussed before, the number of optimal metabolic networks for a given problem can be extremely large, and so classical vertex enumeration techniques are not suitable for this task. One reason is that, given the potential vast number of possible solutions and a fixed amount of time to generate a variety of optimal solutions, there is no guarantee that these methods will generate a diverse set of solutions. In fact, the opposite is more likely: similar solutions (e.g., small variations in reactions on the same pathway) will probably be closer in the search space. Also, due to symmetries introduced by loops and other patterns in metabolic networks, chances are that the enumeration gets trapped performing enumeration in small dense regions of the search space that can be more related to artifacts than to solutions with true biological meaning.

Instead, we advocate for generating a diverse set of solutions, that is, given some experimental condition for which we cannot characterize the metabolic state with just one optimal network, we want to obtain a sample *O* ⊆ Ω^∗^ of this largely unknown set of possible networks in a way that covers well the range of possibilities. In other words: if large differences in metabolism can be explained by the same experimental data, we want to obtain a diverse set of these optimal networks that capture those different metabolic states. This usually means exploring distant solutions with changes that correspond also to distant pathways.

The concept of diversity of optimal solutions of a MILP problem is not well explored in metabolic network reconstruction, and only marginally analyzed in combinatorial optimization. Of special interest is the sequential MILP approach proposed by Danna et al. [32], in which they propose an enumeration strategy which incorporates the concept of diversity by maximizing the distance to previously found solutions at the same time that they discard visited solutions. The closest concept to this general strategy applied to the enumeration of optimal context-specific metabolic networks can be found in [20], where Robaina et al. incorporate the idea of maximizing the distance to the previous solution, but without a mechanism that would remove already explored solutions.

Although maximizing the distance may seem like a good idea a priori, in practice it can lead to oscillations in the search, in which the search process jumps between two distant clusters of possible networks, with large inter-cluster distance but very small intra-cluster distance. That is why the concept of diversity in metabolic networks must be carefully analyzed with synthetic and real data that allow observing the behavior and quality of the solutions. In the following sections we present some enumeration strategies and analyze their advantages and drawbacks. Based on these limitations we introduce a new way to generate diverse solutions that we later evaluate in the context of prediction of essential genes, by constructing ensembles of diverse optimal networks.

It should be noted that we limited to a set of generic techniques that can be implemented on top of general MILP solvers and can be easily integrated in the existent pipelines for network reconstruction. One disadvantage of this is that each solution is obtained by constructing and solving a new MILP problem. Ad-hoc search strategies for the enumeration of MILP solutions based on custom branch-and-cut methods or more advanced tree search exploration, although they might be more efficient in some situations, are out of the scope of this work.

### Enumeration of optimal networks by inclusion or exclusion of reactions (Rxn-enum method)

A simple way to generate alternative optimal metabolic networks can be achieved by directly manipulating the flux bounds of each reaction to force it to carry some positive flux, some negative flux (if reversible), or no flux, as in [18, 19]. The original method traverses all the reactions in the model testing forward (or backward flux if the reaction is reversible) or blocking flux in order to generate a new solution with a different activation for each reaction. Solutions that are still optimal after the modification are added to the set of optimal solutions. This method has however two major limitations: 1) it only explores variations in single reactions (if they can be active or inactive in an optimal solution), leaving the vast space of combinations between reactions completely unexplored; and 2) it generates many duplicated solutions, wasting computation time.

A very naive modification to this basic algorithm to alleviate the second issue consists in tracking the activation or inactivation of each reaction in the set of alternative optimal networks during the search process. If forcing the flux through a reaction *R*_*ijk*_ results in an optimal sub-network with another reaction *R*_*i,j*+1,*k*+1_ active, then there is no need to force flux through *R*_*i,j*+1,*k*+1_ as it is not guaranteed that this operation is going to generate a new sub-network (unless the solver is adjusted to increase randomness in the solutions returned).

One advantage of this approach is that it tests every reaction in the model to see if its presence or absence affects the quality of the solution. This generates alternative networks with modifications in every possible pathway of the metabolic network, which makes this technique a good starting point for more advanced enumeration methods (for example, to generate a set of initial candidate optimal solutions).

### Exhaustive enumeration of optimal networks (integer-cut method)

One simple way to perform a full enumeration of the set of optimal networks is by adding integer-cuts [33]. Starting with a default optimal solution ***x***^∗^ to the MILP problem defined in Eq. 3, a new solution is generated by adding a new constraint to the original problem to cut ***x***^**∗**^ from the set of feasible optimal solutions. This process is repeated for each new solution, adding a new constraint per solution. A new solution is accepted if there is at least one different reaction in the candidate sub-network, that is:

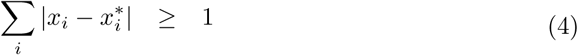

Although this constraint is not linear due to the absolute value, it can be linearized by considering separately the ones from the zeros. Two solutions are equal if they have the same set of active reactions and the same set of inactive reactions. Thus, for each 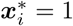, we expect to have *x*_*i*_ = 1, and for each 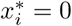, we expect *x*_*i*_ = 0 if both the previous solution and the candidate are equal. Under this situation, summing up all the ones from *x*_*i*_ for which 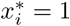 should be equal to 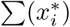 (except if there is one or more differences), and in the same way, summing up all the zeros from *x*_*i*_ for which 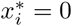 should be equal to zero. If this does not happen, then there is some difference between the candidate solution *x*_*i*_ and a previous optimal solution *x*^∗^. More formally, the linearization of Eq. 4 can be written as:

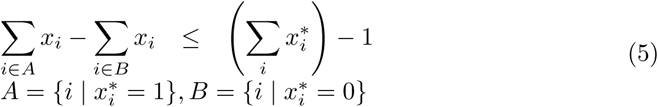

By adding this constraint for each new ***x***^∗^ returned by the solver, we exclude all the previous solutions that have been found so far. The generation of new solutions stops when the problem becomes infeasible, that is, there are no more feasible optimal solutions. Note that this cut can be modified to cut feasible optimal solutions that differ in more than one reactions, i.e., to cut solutions that are at some specific hamming distance.

The advantage of this method is that it enumerates all possible solutions since it removes one by one every optimal feasible solution. It is straightforward to see that this method enumerates all the feasible optimal solutions by observing that: 1) each cut removes one optimal solution; 2) the number of constraints that are added grows monotonically with every new optimal solution; and 3) the number of solutions is finite. Let us assume that for a given problem, the set of optimal feasible solutions Ω^∗^ contains *N* different solutions, i.e., for every pair 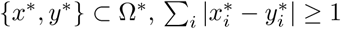 (there is at least one different reaction between any two optimal solutions). For the sake of the proof, let us assume that after *N* steps of the algorithm, and after adding *N* integer cuts, one per optimal solution, the last MILP problem is still feasible, i.e., solving it returns a solution *z*^∗^, thus: 1) *z*^∗^ is different to any other solutions in at least one reaction, which means that there are at least *N* + 1 solutions, contradicting the initial assumption; or 2) *z*^∗^ is a duplicated solution, that is, there exist a solution *x*^∗^ ∈ Ω^∗^ such that 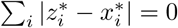, which contradicts the definition of the integer cut.

However, in practice, it is not possible to enumerate the entire space of solutions due to the potential number of possible optimal solutions. Although this technique can be used also to generate a sample of optimal solutions (stopping the search after a desired number of solutions was found), the method is not well suited for this task since: 1) the number of constraints grows linearly with the number of solutions, which increases the difficulty with each new solution; 2) the algorithm can get trapped enumerating solutions in a small region of the whole space of possible optimal solutions, and so diversity in the set of solutions is not guaranteed; 3) even if a new optimal solution exists, due to numerical instabilities or precision errors, the search process can prematurely stop at the first incorrectly detected infeasible problem.

### Enumeration of optimal networks with maximum dissimilarity (Maxdist method)

Another strategy for the enumeration of optimal solutions is to search the most dissimilar metabolic network to a previous optimal one. This idea, already explored in the context of Integer Programming problems [32, 34], has been also proposed for metabolic networks [20]. The strategy requires to solve a bilevel optimization problem in which the inner optimization problem solves the original problem and the outer optimization maximizes dissimilarity. This particular bilevel optimization can be implemented as a standard MILP problem, by introducing a constraint that corresponds to the original objective function. First, an optimal solution ***x***^∗^ with optimal score *f*(***x***^∗^) = *z*^∗^ is calculated using the problem defined in Eq. 3, and then the original objective function is replaced by the minimization of a function *g*(***x, x***^∗^) which measures the similarity between the candidate solution ***x*** and a reference optimal solution ***x***^∗^. In order to guarantee that the new solution to this new problem is still optimal in the original problem, a new constraint *f* (***x***) = ∑_*i*_(*x*_*i*_) = *z*^∗^ has to be added to preserve optimality.

Although the idea of returning the most dissimilar optimal network is interesting, one of the limitations is that it can easily oscillate between a small set of optimal networks that are the most distant to each other, since only the previous optimal solution is discarded. In order to break this oscillatory pattern, we can introduce integer cuts to discard already visited solutions. This modification prevents trivial oscillations between already visited solutions and enumerates the space of solutions starting from the most extreme differences. The objective function *g* can be defined as the minimization of the overlapping of ones between ***x*** and ***x***^**∗**^. Note that the optimality constraint guarantees that the solutions must have the same number of ones (same score), and so removing one overlap (for example by not including a reaction in *R*_*H*_ which is present in the reference solution) has to be compensated by including another reaction in the set of *R*_*H*_ not present in the reference solution, or by removing one reaction in the *R*_*L*_ set which is present in the reference solution, in order to preserve the original optimal score. Minimization of the overlapping of ones between ***x*** and ***x***^∗^ with this constraint is essentially the same as finding the most extreme vertices of the 0/1-polytope of feasible optimal solutions using the hamming distance.

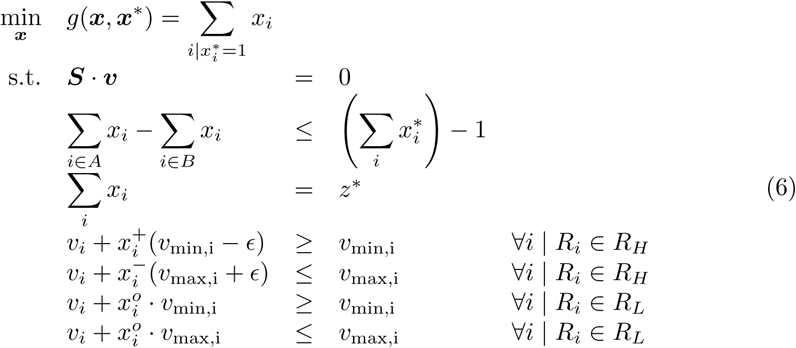

The expected behavior of this algorithm is the following: starting from the default solution ***x***^∗^, the search process generates the most distant network with the same optimal score. This process is repeated, changing the ***x***^∗^ in each iteration to the one previously found, pushing away the search to the boundaries of the space until the most distant networks in the space of optimal solutions are discovered. The `integer-cut` constraint prevents search loops, and so once the extremes are found, the distance of the new discovered solutions decreases progressively.

This method has also limitations that may prevent its use for generating a diverse sample of optimal solutions. Concretely, even though the integer-cut constraint prevents generating repeated solutions, the density of similar metabolic networks at the boundaries can be large enough to never explore other areas. This increases the risk of ending up oscillating between a small group of clusters of networks with a large inter-cluster distance but a very small intra-cluster distance. In addition to this, the method is computationally more expensive than the previous ones.

### Measuring diversity

Given the unknown volume of the 0/1-polytope comprising the optimal solutions, it is not possible to directly estimate its size without sampling solution from it. In order to measure how diverse are the set of solutions obtained with different methods, we need to rely instead on indirect measures. Since solutions are indexed by 0/1 coordinates, one reasonable metric to use is the hamming distance:

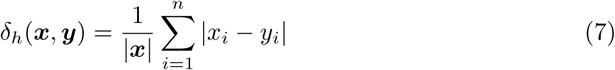

For each pair of solution vectors ***x, y*** ∈ {0, 1}^*n*^ obtained from the set of optimal solutions Ω^∗^, we compute the hamming distance (i.e., how many reactions are different between any two solutions) and we average across all the distances to obtain the *average pairwise distance* 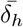. One way to promote diversity is to maximize this measurement: between two different sets of optimal solutions (of a similar size), the set with a larger average pairwise distance contains solutions that are, on average, more diverse.

However, relying only on the average pairwise distance might not be informative enough in some situations. For example, as mentioned before, if the enumerative strategy discovers at the beginning the most two distant solutions, and then enumerates similar solutions to those initial maximal distant solutions, the average pairwise distance is going to be maximal at the beginning of the search. Under these circumstances, it is easy to have the false impression that the set of solutions is diverse, but instead it will contain only the two initial different solutions with very small variations.

To discriminate better between these situations, we use also the *average nearest neighbor distance* 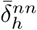 defined as:

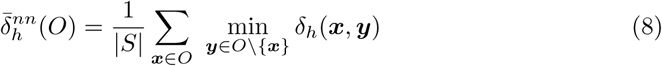

That is, for each optimal solution in the solution set *O* ⊆ Ω^∗^ obtained with some enumeration method, we measure the distance to all other solutions and we take the distance to its closest solution (nearest neighbor). Then, we average all those distances to have the average nearest neighbor distance.

The average nearest neighbor distance measures how *spread* the solutions are. We want solutions that are spread to cover a wider range of different solutions and avoid the enumeration of clusters of very similar solutions.

Considering these two metrics, we can devise four situations when comparing the solution sets obtained by different methods:

- **Lower** 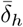 and **lower** 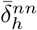: this situation corresponds to a low diversity. Solutions are close together and sampled from a small region of the search space.
- **Larger** 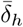 and **lower** 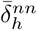: low dispersion of the solutions, even though solutions are distant from each other.
- **Lower** 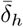 and **larger** 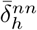: solutions are dispersed but only in a smaller region of the search space.
- **Larger** 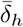 and **larger** 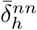: better diverse set of solutions in which solutions are scattered across the space of optimal networks.

Under these terms, we say that one method is more diverse than another if the distance of both measurements is higher for a sample of optimal solutions of predetermined size.

### Diversity based extraction of optimal metabolic networks (DEXOM method)

Based on the previously identified problems and improvements for each method, we propose a new method to generate a set of diverse optimal metabolic networks, combining the advantages of the techniques described before. The basic steps of `DEXOM` are:

1. Generate an initial set of optimal solutions using the `Rxn-enum` method with integer cuts to avoid duplicated solutions.
2. Pick an initial solution ***x***^**(0)**^ from this set.
3. Use the selected solution to create a template vector and to find a solution maximizing the dissimilarity with the given template, using the `Maxdist` method. That is, initialize a vector of zeros as a template and copy *n* random reactions from the selected solution to the template (copy *n* random ones from the selected vector to the template). Maximizing the distance to this template will result in a new solution ***x***^**(1)**^ with maximal distance to the selected reactions. The number of chosen reactions (the step size) depends on the current iteration *i* and a parameters *d*_*s*_. As the search progresses, more reactions are picked from the selected solution and the new generated solutions are increasingly different from each other.
4. Set the new solution ***x***^**(1)**^ as the new initial solution and repeat from 3 until the desired number of solutions has been reached or until there are no more solutions.

**Algorithm 1.**
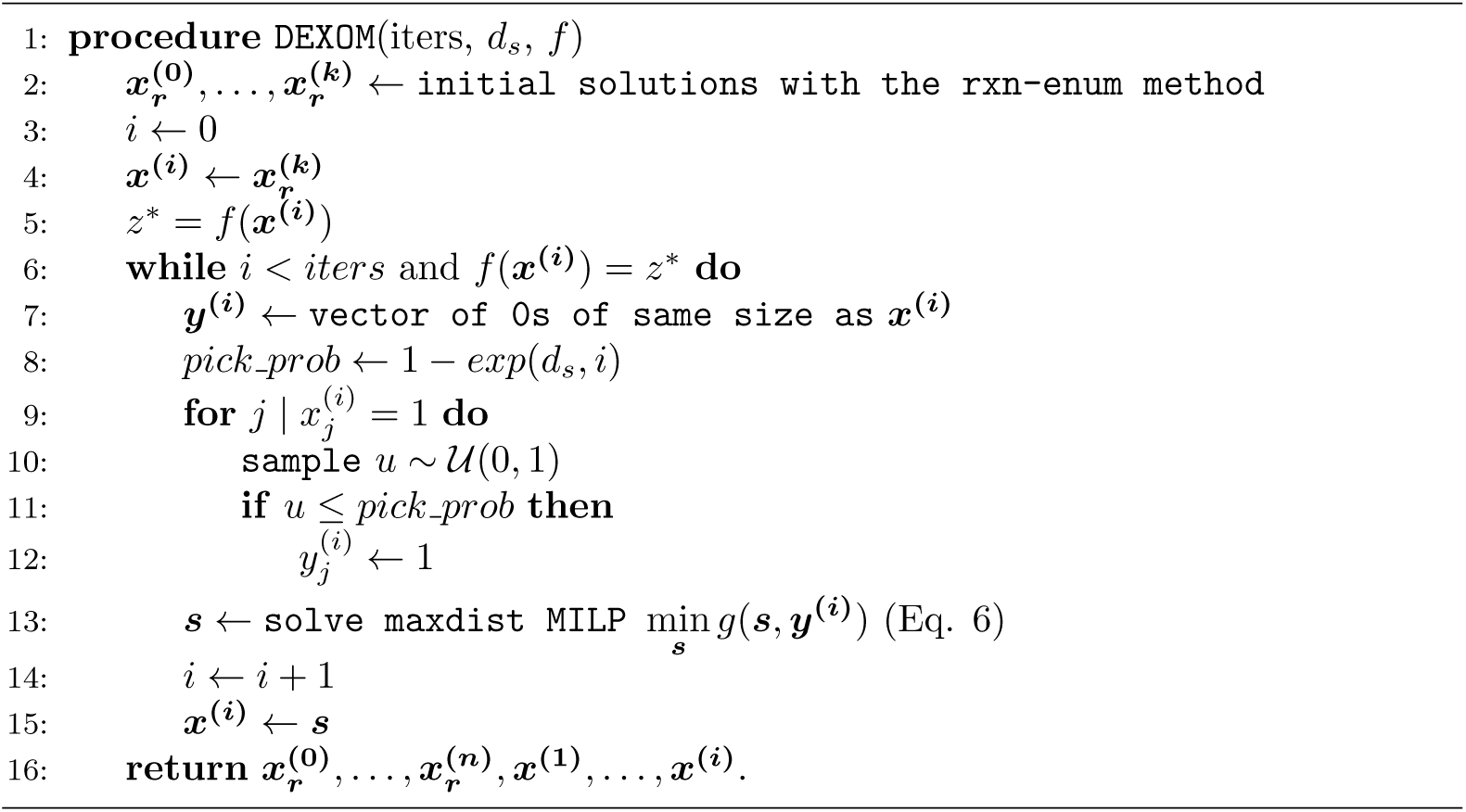
DEXOM algorithm.

A detailed version of the algorithm is described in Alg. 1. `DEXOM` combines the advantages of the previous techniques. It starts computing an initial set of solutions using the Rxn-enum method avoiding duplicated solutions. This guarantees that single variations of reactions across all pathways are explored. Then, starting for any solution from this initial set, the algorithm explores solutions in the vicinity of the selected solution, using it as a *template*. The vicinity distance is controlled by the parameter *d*_*s*_, which controls how fast the step size increases over time. Using a *d*_*s*_ value close to one (e.g. *d*_*s*_ = 0.99), the search concentrates at the beginning with more probability in the close vicinity of the selected solution. This occurs because at the beginning (e.g. iteration *i* = 2), the probability of picking a reaction from the selected solution to the template is *pick_prob* = 1 − *exp*(0.99, 2) = 1 − 0.9801 = 0.0199, which is close to 0, and none or very few reactions are going to be selected. Then, the `Maxdist` method is used to find an alternative solution minimizing the overlapping against the randomly selected reactions in the template ***y***. As the search progresses, the radius of the exploration increases (more reactions are selected, and the distance of the new solution to the selected solution increases), as well as the average distance of the new solutions. As *pick prob* starts to approach to 1, the search process starts to behave more like the `Maxdist` method.

### Essential gene prediction and metabolic network ensembles

Context-specific metabolic networks can be used to make predictions about the metabolism of a cell or tissue in a specific experimental condition. Of a particular interest is the prediction of essential genes. An essential gene is a gene that is indispensable for the organism to survive. Some genes are absolutely required for survival, whereas other genes are conditionally essential, meaning that they are essential depending on the environmental conditions. For example, in *Saccharomyces Cerevisiae*, gene ARG2, which encodes glutamate N-acetyltransferase —a mitochondrial enzyme that catalyzes the first step in the biosynthesis of the arginine— is essential only in the absence of arginine in the medium.

Many essential genes that are related to metabolism (those related to enzymes) can be predicted using metabolic networks. However, conditionally essential genes are particularly hard to predict since they cannot be predicted without integrating experimental data or knowledge related to the condition. Context-specific metabolic networks are able to predict them indirectly, by extracting first the sub-network which is most consistent with the experimental data. After removing all the reactions that are predicted to be inactive in a given context, conditionally essential genes that were not essential in the generic network might be now predicted to be essential.

Predictions of essential genes using metabolic networks can be done by comparing the maximum flux through the biomass reaction —an artificial reaction that encodes the minimum requirements of the organism to sustain a basic metabolic activity— using Flux Balance Analysis (FBA) [35] before and after knocking out a gene in the metabolic network. If the flux through the biomass reactions is below a certain threshold after KO (e.g., below 1% with respect to the wild-type) then the gene is considered essential.

However, as explained before in this section, it is common to find more than one optimal context-specific metabolic network for a given condition, each one representing a slightly different hypothesis of the metabolic state. Each network may predict different essential genes. Since all networks fit the experimental data equally well, there is no clear way to decide a priori which of these predictions may be true. In this situation, a reasonable strategy is to consider that if a network predicts a gene to be essential, then the ensemble decides that the gene is essential, in order to maximize the number of true essential genes (at expenses of increasing the false positives), similar to what has been done in [21] with Gap-Filling methods.

Figure 3 shows an example of how the procedure works. For each gene, a KO is simulated by maximizing the flux through the biomass reaction after knocking out the reaction or reactions associated to the gene (based on the Gene-Protein-Reaction rules), using the `singleGeneDeletion` method from the COBRA Toolbox. If the ratio between the KO and the wild type is below 0.01 (flux after KO below 1%), the gene is classified as essential. This process is repeated for all genes and for all optimal networks, and then results are combined by performing a logical OR of the predictions across networks.

**Fig 3.**
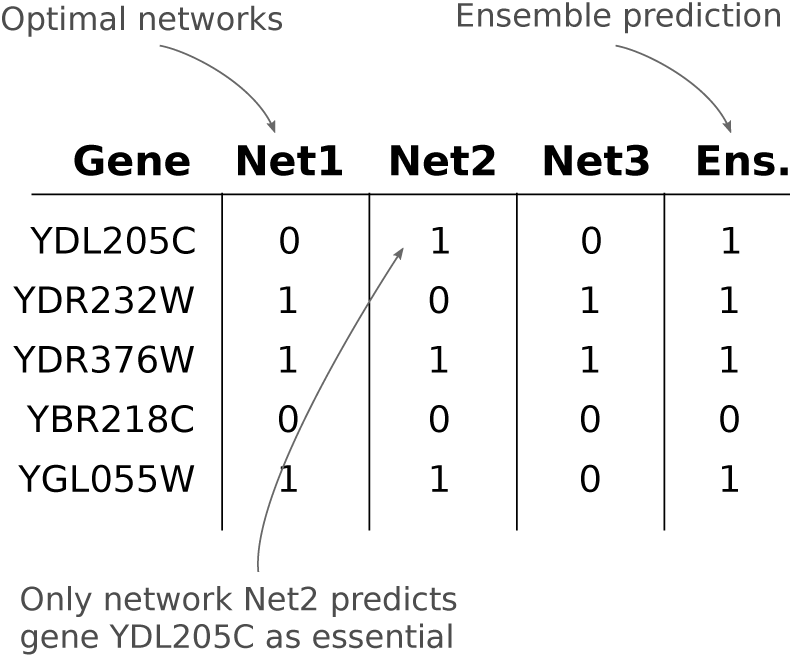
Example of essential gene classification for three optimal networks and the result for the ensemble combining the prediction by performing a logical OR.

After obtaining the predictions for each gene, the True Positive Rate (TPR, sensitivity) and the False Positive Rate (FPR, 1-specificity) are calculated by comparing the predictions against the true essential genes for *Saccharomyces Cerevisiae* (included in the repository of the code), and applying the following formula:

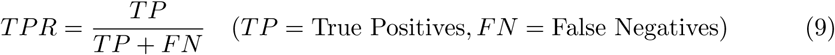

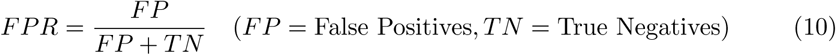

## Results & Discussion

In Section *Methods* we show how the problem of context-specific metabolic network reconstruction is subject to significant variability due to the vast number of possible optimal metabolic networks that explain the same experimental data. This variability makes the interpretation of the metabolism using a single metabolic network not very reliable, since many equally valid alternative hypotheses are disregarded.

In this section, we analyze the performance of each of the proposed methods to generate a diverse sample of optimal metabolic networks, assuming that in practice it is not possible to fully enumerate the total unknown space of optimal solutions, as is generally the case. For this purpose, first we evaluate the diversity of the recovered samples with each method when the true number of optimal solutions is known. We use the DAG model introduced in Section *Methods* as a ground truth generator to construct a problem with a fixed number of possible alternative optimal metabolic networks. Afterwards, we evaluate the different methods in a more realistic setting using the Yeast 6 model. Using this GEM, we select random sets of highly expressed and lowly expressed enzymes to generate problems in which the total number of optimal solutions is not known a priori, and we compare the samples generated with each method in terms of diversity. Finally, we evaluate the predictive capabilities of each method for in-silico prediction of essential genes. Using real gene expression data for *Saccharomyces Cerevisiae* under aerobic conditions [36] and the Yeast 6 model, we enumerate thousands of optimal networks with each method and we asses the performance by predicting which genes are essential using both the individual networks and ensembles of networks constructed by combining the predictions of the individual networks.

### DEXOM improves the diversity of recovered optimal networks

We measure how well each method performs to generate diverse samples of optimal solutions. To do so, we generate samples of fixed size with each method and we measure the diversity of the sample using the average hamming distance and the average nearest neighbor that were introduced in Section *Methods*. We consider two different scenarios: 1) obtaining a sample of optimal metabolic networks in a simulated scenario where the number of total optimal solutions is known; and 2) obtaining a sample of optimal solutions in realistic scenarios where the total number of optimal solutions is unknown.

#### Evaluation in a simulated scenario with a known number of possible optimal solutions

One of the difficulties of measuring the diversity of the solutions obtained by different methods is the absence of a ground truth to compare with, as the full set of optimal solutions is in general not known. However, the DAG network model introduced before can be used as a simple ground truth generator, since the full set of optimal solutions is easy to determine.

In order to assess the coverage and diversity of a sample of optimal networks, we used the DAG network model with 5 layers and 4 metabolites per layer (74 reactions and 22 metabolites in total), which contains a total of 1,024 optimal metabolic networks. The different methods were used to sample from 1 to 250 optimal solutions (around 1/4 of the total set of possible optimal solutions).

Fig. 4 shows a low-dimensional projection of the 250 optimal solutions obtained by each method, where each point is an optimal metabolic network encoded as a binary vector. The grey points correspond to the 1,024 optimal solutions of the ground truth.

**Fig 4.**
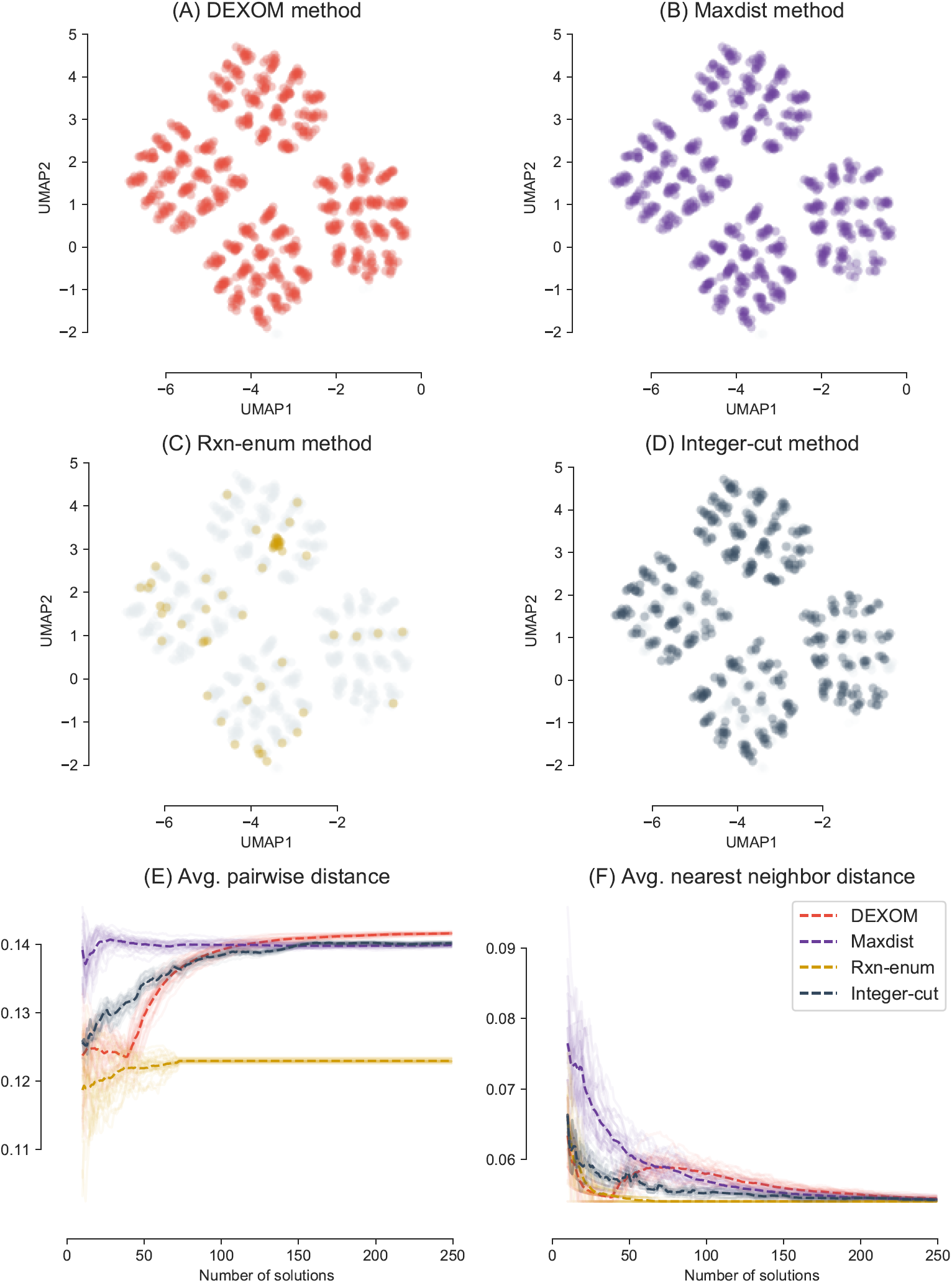
Low dimensional representation of the optimal networks enumerated with different methods (max 250 optimal networks). Each point represents an optimal metabolic network as a binary vector projected in 2D using UMAP with hamming distance and 30 neighbors.

The `Rxn-enum` method shows a low coverage of the space of optimal solutions, enumerating only a 7% of the full space of optimal networks. This is due to the fact that the `Rxn-enum` method changes the bounds of each reaction in the network independently from each other. Since each reaction participates in many optimal solutions, the `Rxn-enum` can obtain only a subset of all possible optimal networks, missing a large fraction of optimal metabolic networks that cannot be recovered with this method.

Qualitatively speaking, the 250 solutions obtained with the `integer-cut` method are not as spread as the ones obtained with `DEXOM` and the `Maxdist` method. Differences between `DEXOM` and `Maxdist` are less obvious and hard to appreciate in a low dimensional embedding. In order to have a better picture of the diversity of the solutions, we calculated the evolution of the distances 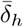 and 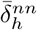 for each method. We repeated the process 30 times to obtain different samples of 250 solutions. The results for the 30 independent runs are shown in Figures 4E and 4F. The average over the 30 runs is represented with a dashed line.

These figures show in a more clear way how `DEXOM` obtains the most diverse set with respect the other methods after 150 optimal solutions were enumerated, surpassing the `Maxdist` method. It can be seen how the behavior of the algorithm in terms of diversity changes dramatically after the initial solution set is calculated (around solution 50). At this point, `DEXOM` starts to increase the distance progressively, looking for more and more distant solutions, which is reflected in the increase of both 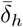 and 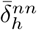. In contrast, `Rxn-enum` obtains sets of solutions with a very poor diversity. After calculating 74 solutions, the method cannot generate new optimal networks (since there are only 74 non reversible reactions in the network), and the solution set stops growing. Since the `Rxn-enum` generates solutions by modifying the constraints of each reaction, one at a time and independently of each other, solutions are mostly concentrated in a concrete region of the space of possible solutions, which corresponds to solutions that are similar to each other. The `Maxdist` method shows at the beginning of the search the largest distance, since the solutions are generated by finding extreme differences. After an initial set of 25 optimal solutions, the average distance stops increasing. This is something to expect since the most distant solutions are usually discovered at the beginning of the search.

#### Evaluation in realistic scenario with an unknown number of optimal solutions

In order to evaluate the diversity in a more biological setting, we randomly select different sets of highly expressed and lowly expressed enzymes of varying size in the Yeast 6 metabolic model [37] and then we enumerate a maximum of 1,000 optimal metabolic networks with the different methods.

Figure 5 shows the results of the enumeration of up to 1,000 optimal sub-networks from a randomly selected set of 120 genes highly expressed and 120 genes lowly expressed on Yeast 6. Enumeration of optimal solutions was repeated 10 times for each method. Since in this case the true set of possible optimal solutions is not known, grey dots represent the union of all discovered optimal networks for all the methods.

**Fig 5.**
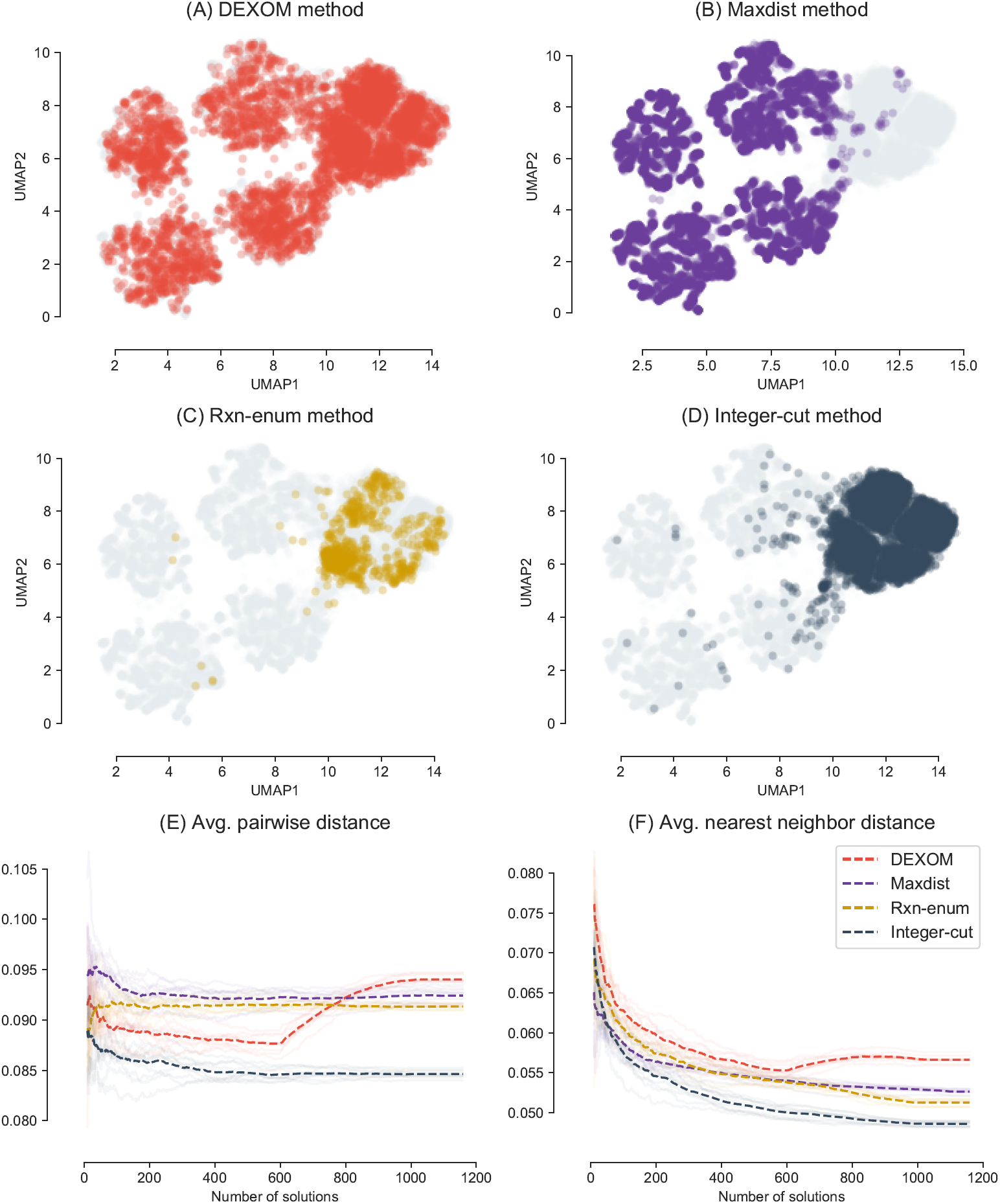
Enumeration of a maximum of 1,000 optimal metabolic networks on Yeast 6 model using a random set of 120 highly expressed genes and 120 lowly expressed genes. Enumeration was repeated 10 times for each method. The grey dots represent the union of all the solutions found by all the methods.

Again, a similar pattern of dispersion of the optimal solutions can be observed as with the DAG model. `DEXOM` (Fig. 5A) obtains a set of solutions that look well spread across the space of enumerated solutions. The `Maxdist` method misses most of the large set of similar solutions that are recovered by the other methods. Both the `Rxn-enum` and the `integer-cut` enumeration explore a similar and restricted region of the space, although `integer-cut` can sample more densely from the same region.

Differences between the methods in this more realistic context are more obvious, and `DEXOM` performs comparatively better than the other methods. After `DEXOM` generates an initial set of around 600 solutions, both the average distance and the average nearest neighbor distance start to grow surpassing the other methods. A similar pattern can be observed for different random sets of selected genes (see Supporting information).

### Prediction of essential genes using ensembles is highly dependent on the strategy used for enumeration

Next, we evaluate the predictive capabilities of the different methods for in-silico prediction of essential genes in the model organism *Saccharomyces Cerevisiae*. We used gene expression measured from yeast in aerobic conditions [36]. Genes were classified into expressed and not expressed using different combinations of thresholds on the quantiles of the distribution as it is commonly done in context-specific network reconstruction. For instance, a threshold of [0.25, 0.75] indicates that genes whose normalized expression value are below the quantile 0.25 are classified as lowly expressed, whereas those above the quantile 0.75 are highly expressed. Reactions were splitted into *R*_*H*_ and *R*_*L*_ sets using the `mapExpressionToReactions` method from the COBRA Toolbox.

Essential genes in Yeast 6 were curated using most updated information from YDPM database and the SGD project [38]. Genes that are essential due to mechanisms not directly related to metabolism were excluded from the set, as they cannot be predicted using FBA. In total, 188 genes out of the 900 in Yeast 6 are considered to be essential under aerobic conditions.

A maximum of 2,000 optimal networks were enumerated for each combination of threshold and method, using a time limit of 8h per threshold/method, and 5 min. timeout for each MILP problem. The lower bound of the biomass reaction was constrained to carry a small positive flux, in order to prevent the generation of networks in which knockouts on essential genes cannot be simulated using FBA. In-silico predictions of essential genes were carried out using COBRA Toolbox v3.0.6, classifying each gene as essential if the flux through the biomass reaction was below 1% after KO.

Essential genes were predicted for each optimal network within the set of the optimal networks obtained by each method and threshold, but also for the ensemble of networks, by taking the union of the predictions as shown in Figure 3. That is, if a gene is predicted as essential by a single optimal network from a set of optimal networks enumerated using a given method and threshold, then the gene is classified as essential by the ensemble. Thus, in total, we generated 16 ensembles per method, one for each threshold.

Table 1 shows the True Positives Rate (TPR, sensitivity) and False Positive Rate (FPR, 1-specificity) of these ensembles. `DEXOM` achieves the best TPR for all thresholds, with the best overall TPR of 0.7713 for the threshold [0.25, 0.90], which corresponds to the correct classification of 145 genes out of the 188 essential genes in the dataset.

**Table 1.** True positive Rate (TPR) and False Positive Rate (FPR) of the ensembles for the prediction of essential genes in Yeast 6, for the different methods and thresholds. Ensembles were generated by taking the union of the predictions of all enumerated networks per method and threshold.

| Threshold | Method | TPR | FPR | Threshold | Method | TPR | FPR |
| --- | --- | --- | --- | --- | --- | --- | --- |
| 0.10 0.90 | Dexom | **0.7234** | 0.1264 | 0.20 0.90 | Dexom | **0.7181** | 0.1194 |
|  | Rxn-enum | 0.7181 | 0.1053 |  | Rxn-enum | 0.7128 | 0.1025 |
|  | Maxdist | 0.4255 | **0.0730** |  | Maxdist | 0.4734 | 0.0969 |
|  | Integer-cut | 0.4681 | 0.0815 |  | Integer-cut | 0.4628 | **0.0576** |
| 0.10 0.85 | Dexom | **0.6755** | 0.0871 | 0.20 0.85 | Dexom | **0.6649** | 0.0927 |
|  | Rxn-enum | 0.6649 | 0.0829 |  | Rxn-enum | **0.6649** | 0.0843 |
|  | Maxdist | 0.4521 | 0.0674 |  | Maxdist | 0.5053 | 0.0702 |
|  | Integer-cut | 0.3617 | **0.0604** |  | Integer-cut | 0.4096 | **0.0365** |
| 0.10 0.80 | Dexom | **0.7128** | 0.0688 | 0.20 0.80 | Dexom | **0.6755** | 0.0520 |
|  | Rxn-enum | **0.7128** | 0.0716 |  | Rxn-enum | 0.6596 | 0.0716 |
|  | Maxdist | 0.4149 | **0.0379** |  | Maxdist | 0.4521 | **0.0337** |
|  | Integer-cut | 0.4096 | 0.0534 |  | Integer-cut | 0.3670 | 0.0562 |
| 0.10 0.75 | Dexom | **0.6649** | 0.0843 | 0.20 0.75 | Dexom | **0.6702** | 0.0590 |
|  | Rxn-enum | **0.6649** | 0.0744 |  | Rxn-enum | **0.6702** | **0.0520** |
|  | Maxdist | 0.4096 | **0.0548** |  | Maxdist | 0.4096 | 0.0632 |
|  | Integer-cut | 0.3723 | **0.0548** |  | Integer-cut | 0.3670 | 0.0534 |
| 0.15 0.90 | Dexom | **0.7287** | 0.1053 | 0.25 0.90 | Dexom | **0.7713** | 0.1334 |
|  | Rxn-enum | 0.7234 | 0.1096 |  | Rxn-enum | 0.7340 | 0.1194 |
|  | Maxdist | 0.4681 | **0.0758** |  | Maxdist | 0.5213 | 0.1067 |
|  | Integer-cut | 0.4628 | 0.0983 |  | Integer-cut | 0.4787 | **0.0913** |
| 0.15 0.85 | Dexom | **0.7128** | 0.0829 | 0.25 0.85 | Dexom | **0.6862** | 0.0885 |
|  | Rxn-enum | **0.7128** | **0.0576** |  | Rxn-enum | 0.6649 | 0.0843 |
|  | Maxdist | 0.5000 | 0.0927 |  | Maxdist | 0.4574 | 0.0730 |
|  | Integer-cut | 0.3617 | **0.0576** |  | Integer-cut | 0.4202 | **0.0590** |
| 0.15 0.80 | Dexom | **0.7021** | 0.0885 | 0.25 0.80 | Dexom | **0.6702** | 0.0871 |
|  | Rxn-enum | **0.7021** | 0.0815 |  | Rxn-enum | **0.6702** | 0.0576 |
|  | Maxdist | 0.3989 | **0.0323** |  | Maxdist | 0.4096 | 0.0590 |
|  | Integer-cut | 0.3670 | 0.0548 |  | Integer-cut | 0.4521 | **0.0534** |
| 0.15 0.75 | Dexom | **0.6649** | 0.0506 | 0.25 0.75 | Dexom | **0.6755** | 0.0843 |
|  | Rxn-enum | **0.6649** | 0.0801 |  | Rxn-enum | 0.6702 | 0.0801 |
|  | Maxdist | 0.4309 | **0.0534** |  | Maxdist | 0.4096 | 0.0604 |
|  | Integer-cut | 0.4043 | 0.0562 |  | Integer-cut | 0.4096 | **0.0548** |

These results are followed by the `Rxn-enum` method, which achieves the same TPR as `DEXOM` in 8 out of 16 tests, with a slightly lower FPR in 6 out of those 8 tests. In contrast, `Maxdist` and `integer-cut` ensembles are not very well positioned in terms of TPR, although both methods achieve the lowest rates of false positives. Concretely, the `integer-cut` method obtained the lowest FPR in 9 out of the 16 tests.

Differences between ensembles can be better assessed by placing each ensemble in a ROC space (Figure 6), in which each point is an ensemble represented by its TPR and FPR. The upper part of the figure is dominated by `DEXOM` and the `Rxn-enum` method, whereas the `Maxdist` and `integer-cut` ensembles are characterized by a lower ratio of true and false positives.

**Fig 6.**
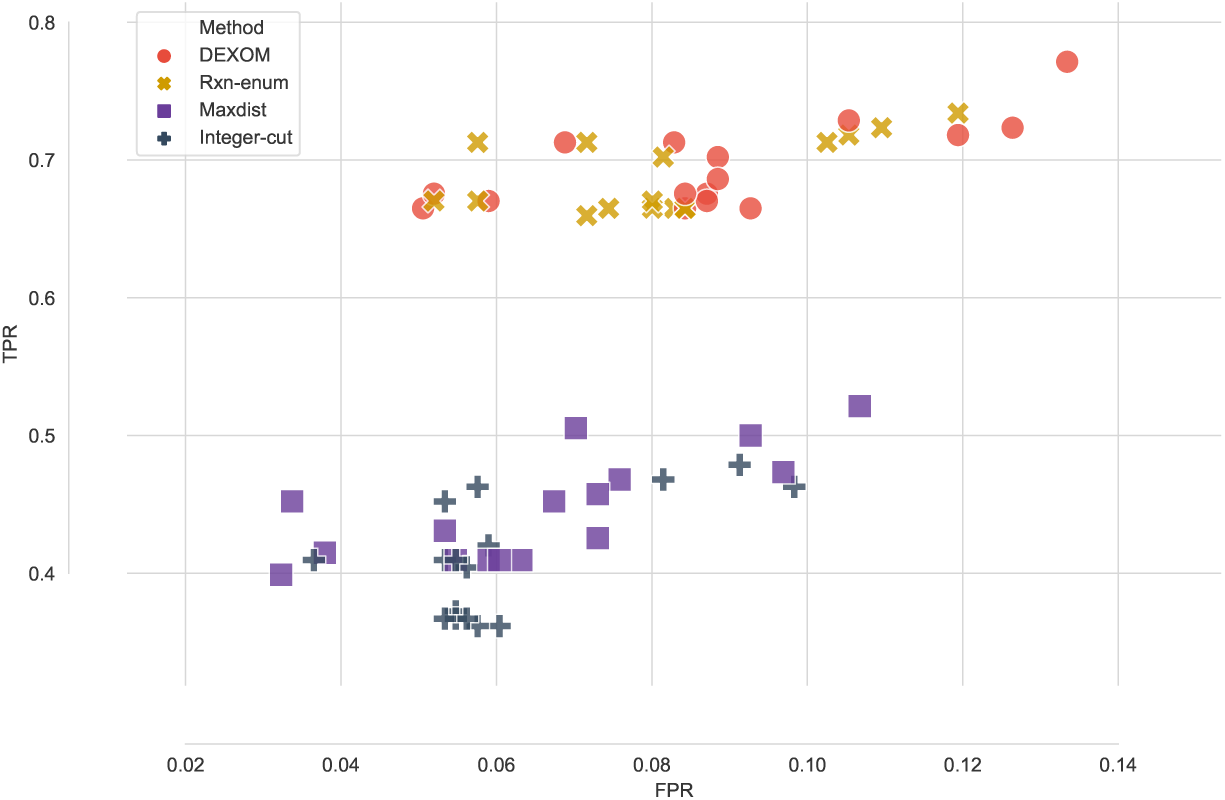
Ensembles of optimal metabolic networks in the ROC space. Each point represents an ensemble of networks built using a method and a concrete threshold for gene expression.

One reason that explains these differences between the methods is the systematic generation of alternative solutions by testing every reaction in the model. If one reaction associated to a gene that is essential is not present in any of the set of optimal networks, the gene is not predicted to be essential. However, if there exist at least one optimal solution in which this reaction is present and essential, both `Rxn-enum` and `DEXOM` have more chances to detect it as they are going to test if there exist an optimal network with that reaction being active. `Maxdist` and `integer-cut` methods leave many of these solutions unexplored. `DEXOM`, in contrast, uses the `Rxn-enum` strategy to have an initial set of solutions with variations in single reactions, from which it expands the search incrementally, increasing the chances of detecting even more essential genes.

Differences in TPR and FPR for the ensembles and the individual networks are shown in Figures 7 and 8 respectively. One interesting observation is that the individual networks generated by the different methods achieve a similar rate of true positives and false positives, and so the higher rates scored by the ensembles using `DEXOM` and `Rxn-enum` are driven by a more diverse set of predicted essential genes. That is, individual networks enumerated by these methods are able to correctly predict different sets of true essential genes, and so the union of those predictions include a more diverse set of detected essential genes. Concretely, the median TPR for the ensembles generated with `DEXOM` and `Rxn-enum` increase 142% with respect the median TPR of their individual networks, whereas the TPR of the ensembles built with `Maxdist` and `integer-cut` increase only 54% and 51% respectively.

**Fig 7.**
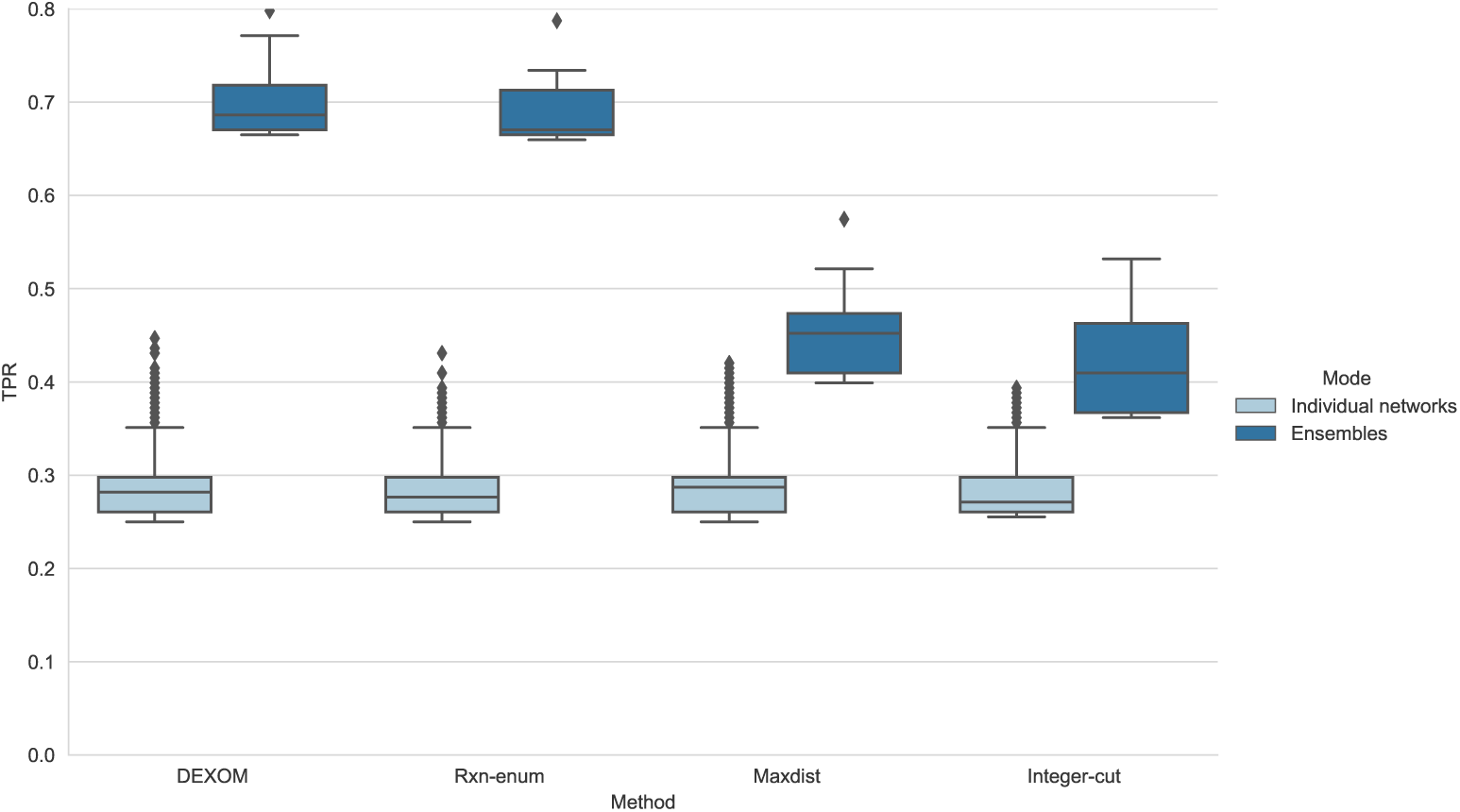
Distribution of the TPR achieved by the individual networks and the ensembles for each method.

**Fig 8.**
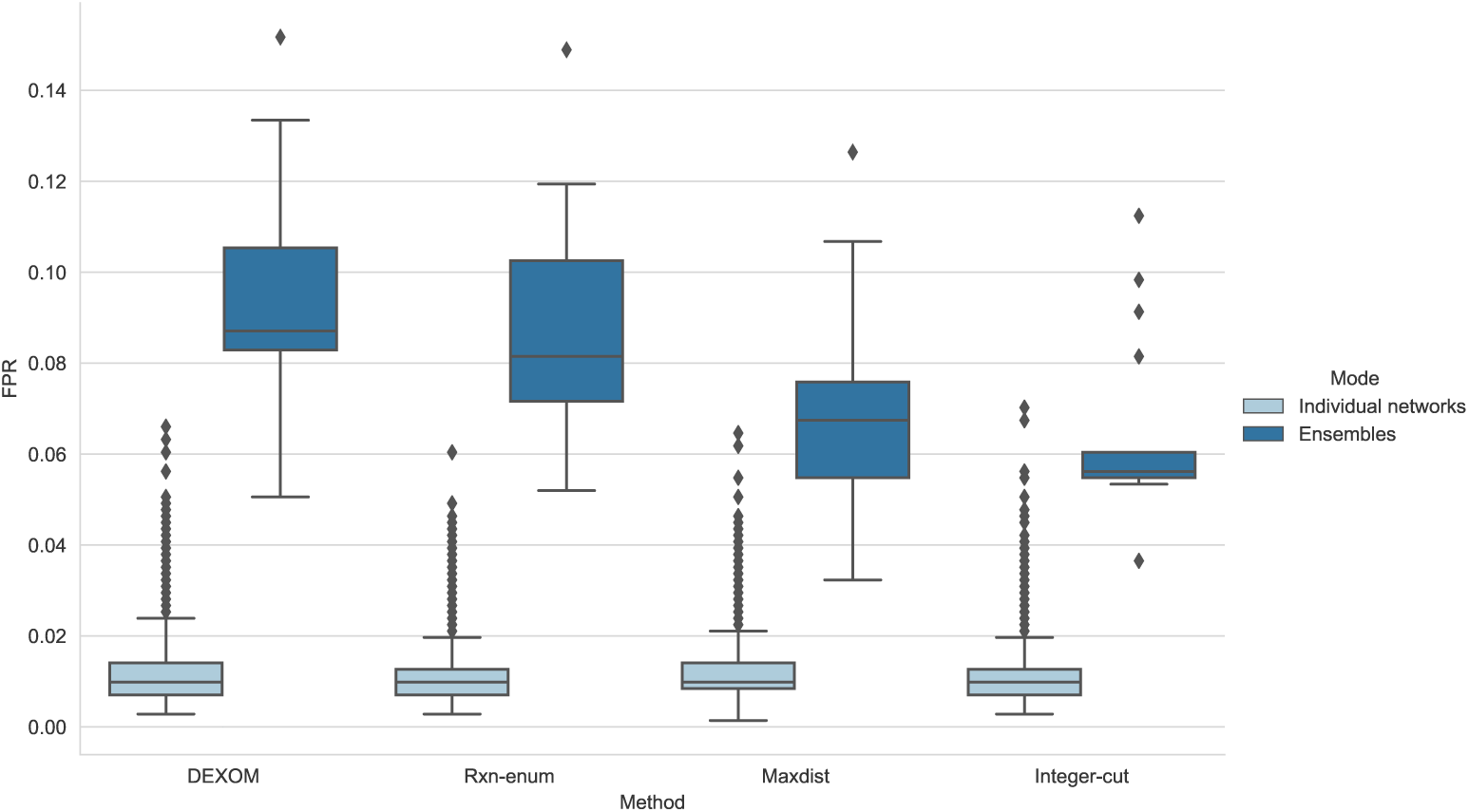
Distribution of the FPR achieved by the individual networks and the ensembles for each method.

The computational cost of the methods is also different. One factor that affects the performance is the number of variables of the MILP problem, and this also depends on the threshold selected. For example, if the number of genes is 1,000 and the threshold for the lowly expressed genes and highly expressed genes is [0.10, 0.90], then only 20% of the genes (200 genes) are used, whereas if the threshold is [0.25, 0.75], 50% of the genes are used. Mapping a bigger set of genes into the networks will translate into larger sets of *R*_*H*_ reactions to maximize and *R*_*L*_ reactions to minimize, and thus bigger MILP problems with more binary variables to optimize.

Figure 9 shows the number of solutions over time for the thresholds used in the evaluation that take the minimum number of genes ([0.10, 0.90], 20% of genes) and the maximum number of genes ([0.25, 0.75], 50% of genes). Figure 9A shows the solutions obtained over time for the case when only 20% of the genes is classified as highly expressed or lowly expressed. In this situation, `DEXOM`, `Maxdist` and `integer-cut` follow a similar trend. `Rxn-enum` is the fastest method, taking less than 10 minutes to finish the enumeration. This is due to the fact that solving each MILP problem has almost no extra cost with respect to the original problem, since only a single constraint to force the inclusion or removal of one reactions is added to the original problem. In contrast, the other methods solve a more complex optimization problem, and the number of constraints grows monotonically, making more difficult the enumeration over time.

**Fig 9.**
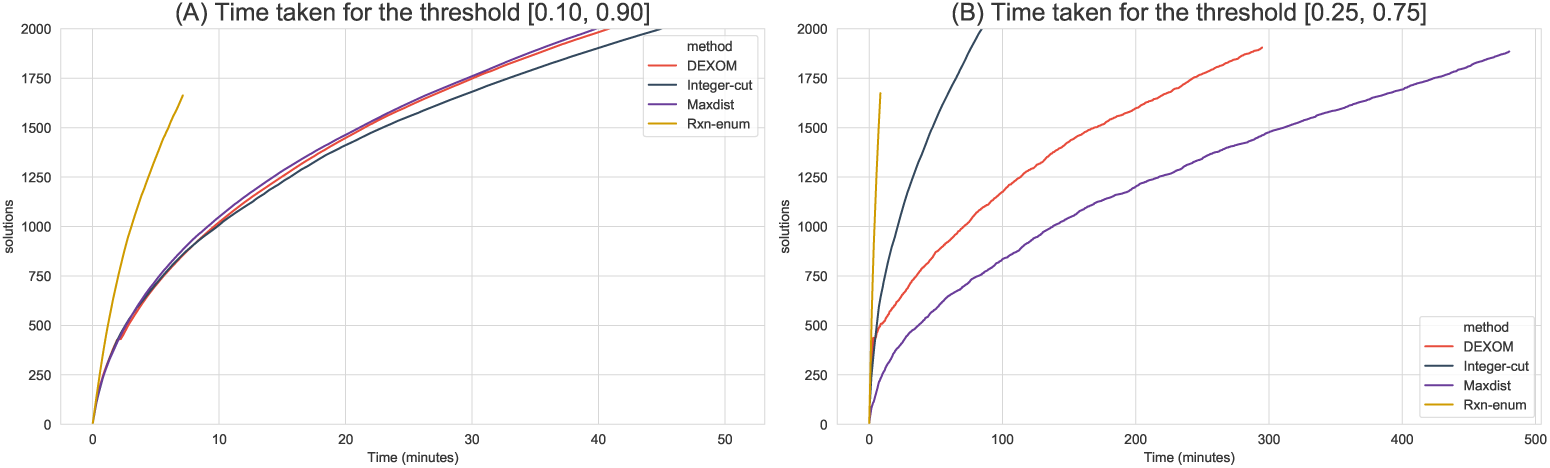
Total number of solutions enumerated over time for the different techniques for two different thresholds. A) Using a gene threshold of [0.10, 0.90]. B) Using a gene threshold of [0.25, 0.75].

Differences between the techniques become more extreme as the number of genes increases. Figure 9B shows the but for the threshold [0.25, 0.75], in which 50% of genes are mapped in the networks. `Rxn-enum` is again the fastest method, followed by `integer-cut`, `DEXOM` and `Maxdist`. Concretely, `Rxn-enum` takes again less than 10 minutes to complete, whereas `integer-cut` takes around 1.5h, `DEXOM` 5h and `Maxdist` around 8h. `Maxdist` is heavily penalized by the increase in the number of selected genes. This is due to the fact that `Maxdist` searches for the most distant vector at each step, and the dimension of this vector correspond to the number of reactions in the *R*_*H*_ and *R*_*L*_ sets, which are bigger in this case. `DEXOM` is less penalized since at the beginning of the search, only a few components of the vector are used to maximize the distance. However, the performance degrades as the distance increases over time, until the distance is maximal. At this point, the performance of `DEXOM` is similar to `Maxdist`. The total time for the evaluation, including all methods and calculation of in-silico predictions of essential gene for each optimal network took around 150 hours in an Intel Core i7 4790 @ 3.60 GHz (8 threads) with CPLEX 12.8 and Matlab 2015 academic.

One important limitation of enumerating optimal solutions is the heavy computational cost involved in the search process. If the number of highly expressed genes and lowly expressed genes is very large, obtaining a single optimal metabolic network can be computational demanding or even not feasible in reasonable time, since obtaining an optimal solution involves solving a MILP problem, which is in general NP-Hard. In this context, enumerating multiple optimal solutions can be prohibitively expensive in some cases, especially with techniques like `Maxdist` or `DEXOM`. One thing that can be done in these situations to alleviate the computational burden is to reduce the integer optimality tolerance of the solver to stop looking for better solutions once the solver has found a feasible integer solution proved to be, for example, within 1% of optimal.

## Conclusion

Context-specific metabolic network reconstruction is a widely used approach to integrate different layers of experimental data into metabolic networks. This process allows to capture the metabolic sub-network that corresponds to the active part of the metabolism of an organism in a given condition. Using this reconstructed model, more advanced techniques such as Flux Balance Analysis, Pathway Enrichment, Network Visualization or Gene Essentiality Prediction can be used to get an integrated view of the metabolic behavior.

One important limitation with this methodology is that context-specific metabolic network reconstruction is subject to significant variability due to the large number of optimal metabolic networks that can be reconstructed for the same experimental data, among other factors. This variability, which is commonly neglected, can contain relevant information and can offer alternative hypothesis of the metabolic state in terms of different combinations of reactions that are predicted to be active or inactive. Thus, the report of results using only a single optimal context-specific metabolic network can be highly biased and can overlook information relevant to the experiment. While this is an important issue, the analysis of the alternative set metabolic networks is a topic not well explored.

In this study we analyze the problem of enumeration of multiple optimal context-specific metabolic networks both from a theoretical and practical perspective. We show how it is common to have multiple different context-specific metabolic networks that optimally explain the same observed experimental data. The set of optimal solutions constitute different hypotheses of the metabolic state and therefore must be taken into account to reduce bias in the interpretation of results.

We analyze the advantages and disadvantages of different methods that can be used for enumerating alternative optimal networks and we introduce `DEXOM`, a novel method for diversity-based enumeration of context-specific metabolic networks. Instead of randomly enumerating optimal networks, `DEXOM` focuses on sampling optimal solutions that are as representative as possible of the space of the unknown possible optimal solutions.

We evaluate the methods both with simulated and real data based on two criteria: 1) diversity of the optimal solutions obtained with each method, using two different distance metrics to evaluate diversity; and 2) predictive capabilities of metabolic networks and metabolic network ensembles generated with each method for the prediction of essential genes in *Saccharomyces Cerevisiae*.

In terms of diversity, `DEXOM` is the method that recovers a more widespread set of optimal solutions, capturing a more diverse set of different reactions among the metabolic networks but equally consistent with the experimental data.

With respect to predictive capabilities of essential genes using the Yeast 6 model, on an individual basis there are no noticeable differences in terms of True Positive Rate (TPR) and False Positive Rate (FPR) between the individual optimal metabolic networks obtained by each method. However, when the results are combined using ensembles of optimal metabolic networks, the TPR of the ensemble obtained with `DEXOM` increases by 140% compared to the median TPR of the individual networks, whereas ensembles generated with the methods that generate less diverse sets of solutions achieved only an increment of 50%. `DEXOM` was also the method the best overall TPR of 0.7713, which corresponds to 145 out of 188 correctly classified essential genes, for a FPR of 0.1334 (95 false positives out of 712 non essential genes). These differences are explained by a more diverse set of essential genes captured by the individual optimal networks enumerated with `DEXOM`.

Overall, this work provides a better method to enumerate optimal context-specific metabolic networks, and highlights the importance of analyzing the space of optimal metabolic networks in a diverse manner, in order to capture as much as possible the variability that is inherent in the reconstruction process.

## Supporting information

**S1 Fig. Enumeration in Yeast 6 with 200 random genes.** Enumeration of a maximum of 1,000 optimal metabolic networks on Yeast 6 model using a random set of 100 highly expressed genes and 100 lowly expressed genes.

**S2 Fig. Enumeration in Yeast 6 with 160 random genes.** Enumeration of a maximum of 1,000 optimal metabolic networks on Yeast 6 model using a random set of 80 highly expressed genes and 80 lowly expressed genes.

**S3 Fig. Enumeration in Yeast 6 with 120 random genes.** Enumeration of a maximum of 1,000 optimal metabolic networks on Yeast 6 model using a random set of 60 highly expressed genes and 60 lowly expressed genes.

## Acknowledgments

This work was supported by grants from the Institut National contre le Cancer (INCa-PLBIO) and the Labex EpiGenMed (“Investissements d’avenir” program, reference ANR-10-LABX-12-01).

---

**S1 Fig.**
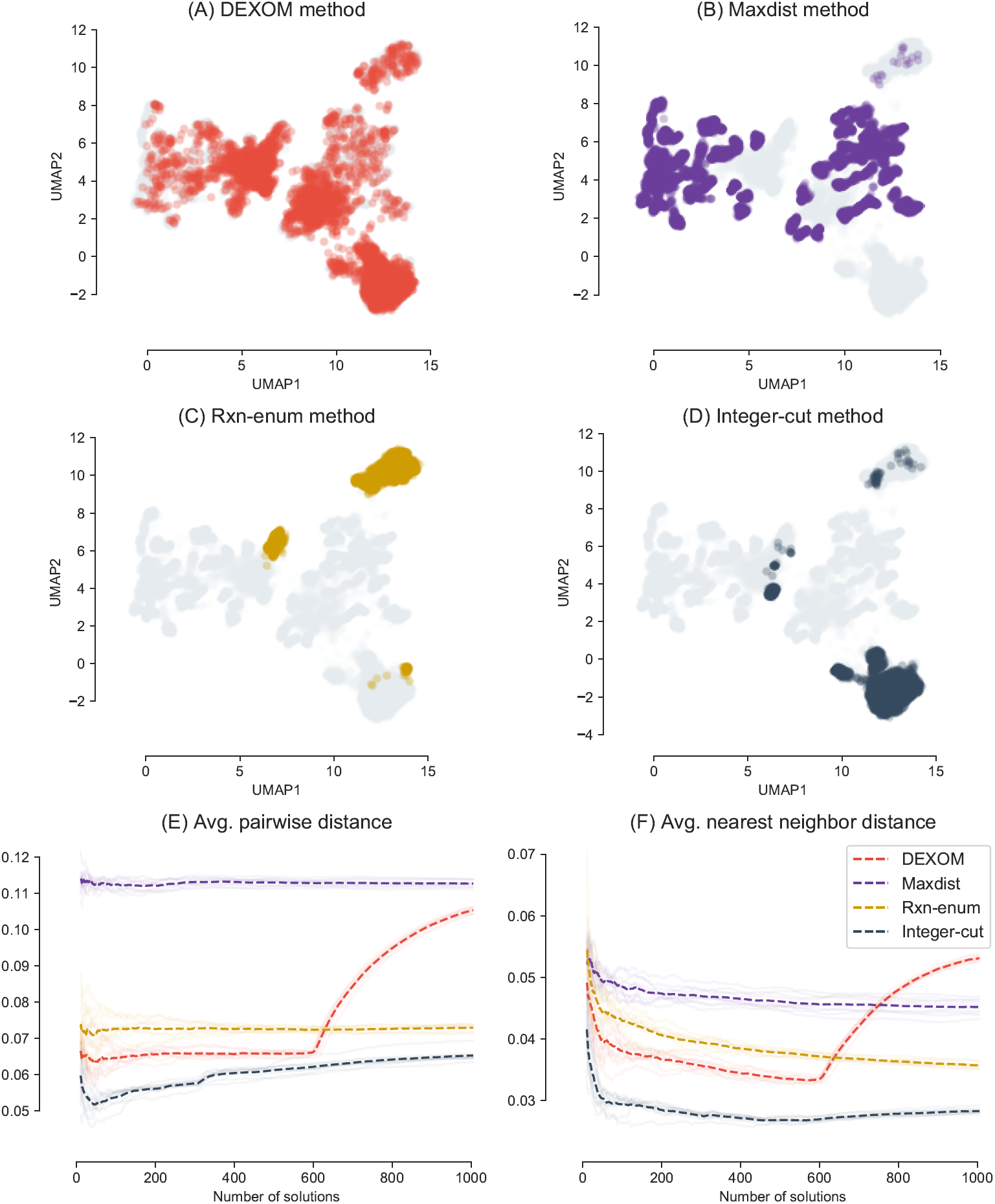
Enumeration of a maximum of 1,000 optimal metabolic networks on Yeast 6 model using a random set of 100 highly expressed genes and 100 lowly expressed genes.

**S2 Fig.**
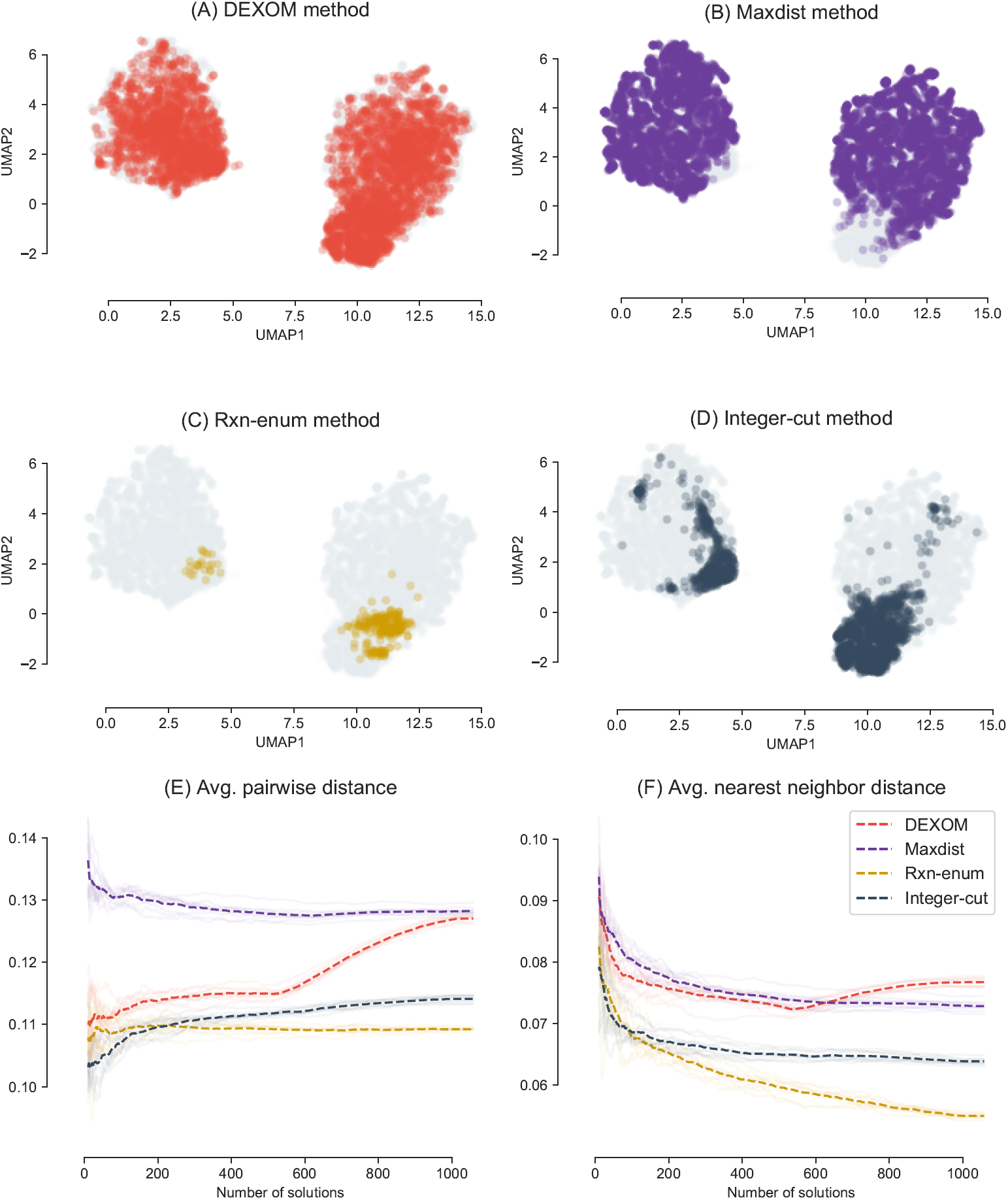
Enumeration of a maximum of 1,000 optimal metabolic networks on Yeast 6 model using a random set of 80 highly expressed genes and 80 lowly expressed genes.

**S3 Fig.**
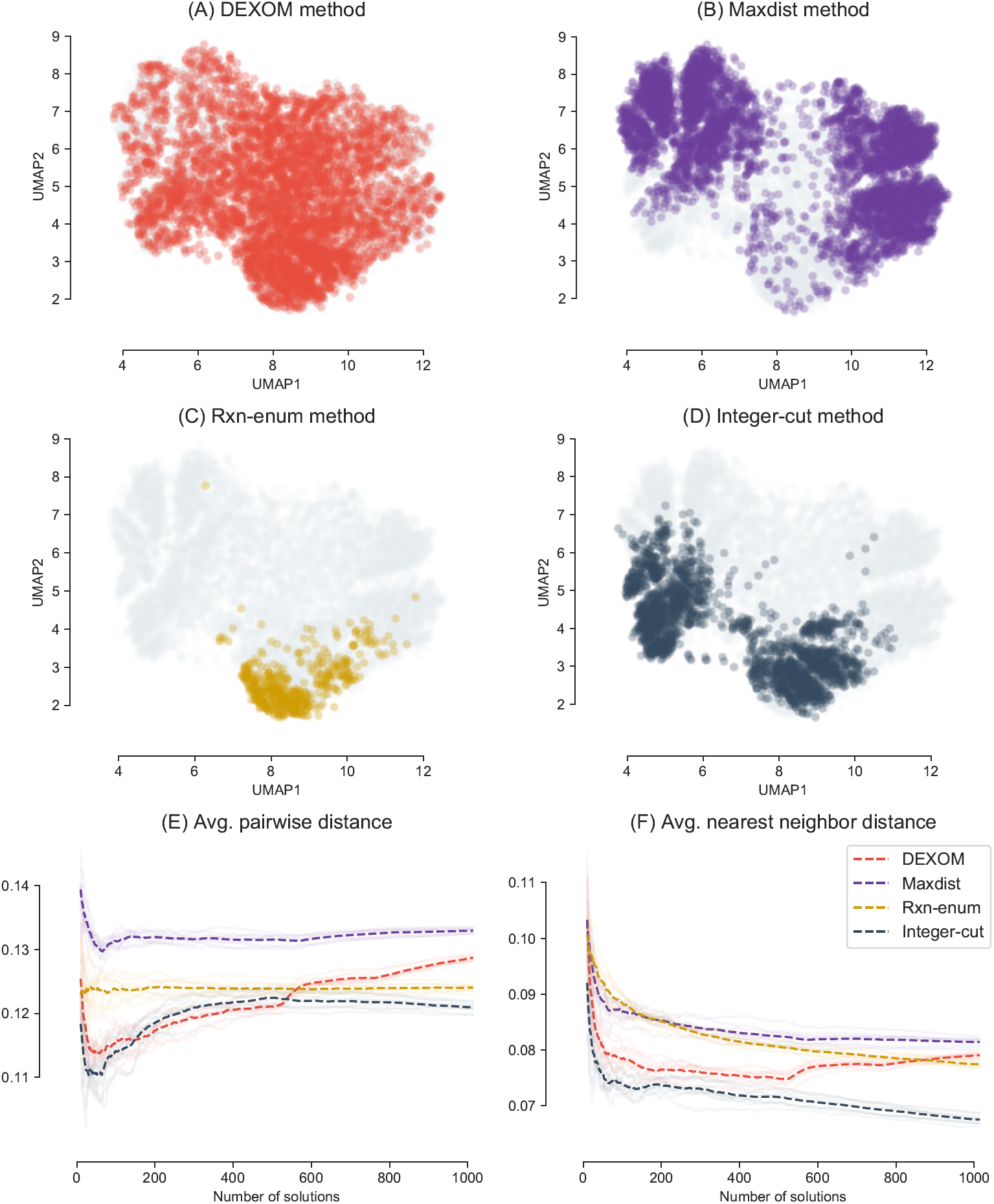
Enumeration of a maximum of 1,000 optimal metabolic networks on Yeast 6 model using a random set of 60 highly expressed genes and 60 lowly expressed genes.

## Footnotes

1 https://github.com/MetExplore/dexom

